# Caspases from Scleractinian Coral Show Unique Regulatory Features

**DOI:** 10.1101/2020.04.13.039685

**Authors:** Suman Shrestha, Jessica Tung, Robert D. Grinshpon, Paul Swartz, Paul T. Hamilton, Bradford Dimos, Laura Mydlarz, A. Clay Clark

## Abstract

Diseases affecting coral have led to massive decline and altered the community structure of reefs. In response to immune challenges, cnidaria activate apoptotic or autophagic pathways, and the particular pathway correlates with disease sensitivity (apoptosis) or resistance (autophagy). Although cnidaria contain complex apoptotic signaling pathways, similar to those in vertebrates, the mechanisms leading to cell death are largely unexplored. We identified and characterized two caspases each from *Orbicella faveolata*, a disease-sensitive stony coral, and *Porites astreoides*, a disease-resistant stony coral. The four caspases are predicted homologs of human caspases-3 and −7, but OfCasp3a and PaCasp7a contain an amino-terminal caspase activation and recruitment domain (CARD) similar to human initiator/inflammatory caspases. In contrast, OfCasp3b and PaCasp3 have short pro-domains, like human effector caspases. We show that OfCasp3a and PaCasp7a are DxxDases, like human caspases-3 and −7, while OfCasp3b and PaCasp3 are more similar to human caspase-6, with VxxDase activity. Our biochemical analyses suggest a mechanism in coral in which the CARD-containing DxxDase is activated on death platforms, but the protease does not directly activate the VxxDase. We also report the first X-ray crystal structure of a coral caspase, that of PaCasp7a determined at 1.57Å resolution. The structure reveals overall conservation of the caspase-hemoglobinase fold in coral as well as an N-terminal peptide bound near the active site that may serve as a regulatory exosite. The binding pocket has been observed in initiator caspases of other species, suggesting mechanisms for the evolution of substrate selection while maintaining common activation mechanisms of CARD-mediated dimerization.

## INTRODUCTION

Apoptotic cell death is thought to be a unique characteristic of metazoans, although its evolutionary origins are unclear. While caspases from human cells, and model organisms such as *C. elegans* and *Drosophila*, have been well-studied both biochemically and structurally (1–6), little is known about caspase activity and regulation from other species (7). Invertebrate caspases were first characterized in *C. elegans* (3, 6) and *Drosophila* (8), but they have proven to be poor models for studying the evolution of the vertebrate apoptotic network as the networks in *C. elegans* and in *Drosophila* utilize fewer caspases and regulatory proteins compared to higher eukaryotes. In contrast, vertebrates have retained many characteristics of the apoptotic machinery found in sponges, sea anemone, and coral (9–11). Genomic studies of cnidarians, the sister group to the bilateria, revealed many genes that were previously thought to have been vertebrate innovations, demonstrating that the extensive gene loss in *C. elegans* and in *Drosophila* resulted in apoptotic pathways that do not reflect the characteristics of ancestral metazoans (12). *C. elegans*, for example, utilizes only one effector caspase (CED-3), which also bears a CARD-motif necessary for its activation (13). In contrast, humans have multiple caspases with discrete functions in the inflammatory and apoptotic pathways (14). Moreover, cytochrome C is not involved in the formation of the apoptosome in *Drosophila*, indicating that this organism lacks the intrinsic pathway found in humans (2). The limitations of these model organisms show that studies of basal metazoans, which appear to have a full complement of apoptotic signaling molecules, are more relevant to the evolutionary pathways of vertebrate apoptotic networks.

Disease susceptibility is one of several major stressors of coral communities, with over thirty-five coral diseases reported that affect over eighty coral species (8, 15). Coral possess a rudimentary immune system that consists of innate immune pathways but no adaptive immune system (16). The invertebrate innate immune system is similar to that of vertebrates in utilizing physical and chemical barriers, cellular defenses, and humoral responses to pathogens (17), but in the relatively new field of ecological immunity, major knowledge gaps remain regarding the cellular defenses to disease (18). Although general response types have been outlined regarding receptor recognition, signaling pathways, and effector responses, such as metabolic changes, very few functional studies have been performed on the responses of coral to disease stressors (19).

Relatively more is known regarding coral responses to temperature stress, and the phenomenon known as bleaching, but there still remain large gaps in our understanding of cellular responses to stress. For example, coral activate cell death responses following expulsion of the algal symbiont (20–23), but cnidarian caspases and their regulation during stress responses have not been studied. In *Aiptasia pallida*, elevated temperatures were shown to induce early-onset of apoptosis in the endoderm where algal symbionts reside (22). Heat-induced apoptosis has also been correlated with the upregulation of the anti-apoptotic protein Bcl-2 in *Acropora millepora*, suggesting that coral possess regulatory mechanisms to compensate for sudden environmental changes (24). Caspase-3-like activity was detected in the stony coral *Pocillopora damicornis* when exposed to heat-stress and high levels of ammonium (25), and caspase inhibitors have been shown to prevent the death of bleached coral (9). Recently, Fuess and colleagues showed an increase in expression of apoptosis-related genes, among others, in Caribbean coral with Eunicea Black Disease, which results in a heavily melanized appearance of gorgonian corals (18), but the response has not been elucidated in functional or biochemical studies. Collectively, the data show the potential for complex apoptotic signaling pathways in coral, but data on activation and control mechanisms, and how they compare to those in vertebrates, are lacking due to a dearth of biochemical characterization.

In order to examine caspase-3-like proteins in coral, we expressed and characterized two caspases each from Caribbean reef-building corals, *Porites astreoides* and from *Orbicella faveolata*. The two coral species are found on opposite ends of the stress-tolerance spectrum, and cellular mechanisms that are activated following an immune challenge correlate to disease-sensitivity (20). For example, *O. faveolata*, a disease-sensitive species, activates caspase-mediated apoptotic pathways upon immune challenge, whereas *P. astreoides*, a disease-tolerant species, activates an adaptive autophagic response (20). These findings suggest that the downregulation of apoptotic genes increases stress tolerance, and upregulation of apoptotic genes exacerbates stress sensitivity. Two of the proteins (called PaCasp7a and OfCasp3a) contain CARD motifs at the N-terminus, an unusual combination that has not been observed in caspases-3 or −7 enzymes from higher eukaryotes. In contrast, PaCasp3 and OfCasp3b show canonical caspase-3/-7 structural organization, with short pro-domains. We describe the first biochemical characterization of the coral caspases and show that the PaCasp3 and OfCasp3b enzymes are not activated directly by the CARD-containing PaCasp7a and OfCasp3a, respectively. We also report the first X-ray crystal structure of a coral caspase, that of PaCasp7a determined at 1.57Å resolution, which reveals an N-terminal peptide bound near the active site that may serve as a regulatory exosite.

## MATERIAL AND METHODS

### Cloning, Protein Expression and Protein Purification

The codon optimized sequences of the four coral caspases, PaCasp3, PaCasp7a, OfCasp3a and OfCasp3b, were based on the sequences from previous transcriptomic data (20) and were cloned into pET11a vector (Genescript, USA). All proteins contained a C-terminal His_6_ tag and were expressed in *E. coli* BL21(DE3) pLysS cells and purified as previously described (26, 27).

### Phylogenetic Analysis

Caspase sequences of representative species were obtained from the CaspBase (caspbase.org) (28) along with BLAST top hits from HMMER (29), and multiple sequence alignments were obtained using MEGA 7 (30). The best model of evolution to construct a phylogenetic tree from our dataset was determined with ProtTest 3 (31) (https://github.com/ddarriba/prottest3), and the tree was computed with the maximum likelihood method in IQTREE, using the Jones-Taylor Thornton model (JTT) plus gamma distribution (32). The tree was bootstrapped 1000 times as a test of phylogeny. The accession numbers of all genes used for phylogenetic analysis are listed in Supplementary Tables S1 and S2.

### Size Exclusion Chromatography

Proteins were examined using a Superdex75 Increase 10/300GL column on an AKTA-FPLC. The proteins were concentrated to 1-5 mg/mL and dialyzed in a buffer of 30 mM potassium phosphate, pH 7.5, containing 1 mM DTT for 4 hours. The column was equilibrated with two columns volume (50 mL) of the same buffer. Protein (200 μL) was loaded onto the column, and the column was resolved at a flow rate of 0.5 mL/min. The column was calibrated using the gel filtration LMW calibration kit (GE Health Sciences) following the manufacturer instructions.

### Mass Spectrometry

Matrix-assisted laser desorption/ionization (MALDI) analysis was done as described (33). In brief, proteins were resolved by SDS-PAGE on a 12.5% acrylamide gel, and then bands for the large and small subunits were excised. Each gel fragment was destained using a solution of acetonitrile and 50 mM ammonium bicarbonate (1:1 v/v) for 3 hrs. The gel fragments were then crushed in microcentrifuge tubes, and the proteins were extracted with 30 μL of a solution of formic acid/water/2-propanol (1:3:2 v/v/v) (FWI) for 8 hours at room temperature. After extraction, samples were centrifuged and supernatant was lyophilized then re-dissolved in 2 μL of MALDI matrix solution (FWI saturated with 4-hydroxy-*α*-cyano-cinnamic acid (4HCCA)). Dissolved protein was then retrieved for MS analysis using dried-drop method of matrix crystallization then analyzed by MALDI-MS (Axima Assurance Linear MALDI TOF).

### Whole-Protein Cleavage Assay

Enzyme specificity of the four coral caspases was first examined by cleavage of human procaspases-3 and −6 in time-course assays, as described previously (34). The procaspase substrate was diluted to a final concentration of 5 μM in a buffer of 150 mM Tris-HCl, pH 7.5, 50 mM NaCl, 1% sucrose, and 10 mM DTT at 37 °C. Reactions were started by the addition of respective coral caspase at a final concentration of 1 μM, and the total reaction volume was 2 mL. Aliquots of 100 μL were removed at times 30 sec, 1 min, 5 min, 15 min, 30 min, 45 min, 1 hour, 2 hour, 4 hour, 6 hour and 8 hour after the addition of active enzyme. Reactions were stopped by the addition of six-fold concentrated SDS-PAGE loading dye (20 μL) followed by incubation in boiling water for 5 minutes. Samples were loaded into a 16% resolving gel with a 4% stacking gel and electrophoresed for 1.5 hours at 80 volts followed by an increase in voltage to 190 V for an additional 4 hours. The change in density for the procaspase substrate over time as a result of cleavage was quantified using Image lab software (Bio-Rad), and the data were plotted with Kaleidagraph. As described previously (34), the data were fit to an exponential decay to determine the CF50 (cleavage of 50% of protein substrate), and the CF50 was used to calculate the rate of hydrolysis (M^-1^ sec^-1^) using the equation k = ((−ln(P))/(E·t)). In this case, k is the rate of hydrolysis, P is the fraction cleaved (50%), E is the concentration at which CF50 is achieved (in molar), and t represents time (in seconds).

### Enzyme Assays and Substrate-Phage Display

Enzyme activity was determined in a buffer of 150 mM Tris-HCl, pH 7.5, 50 mM NaCl, 10 mM DTT, 1% sucrose, 0.1% CHAPS (assay buffer) at 25 °C, as previously described (35, 36). The total reaction volume was 200 µL, and the final enzyme concentration was 10 nM. Following the addition of substrate (Ac-DEVD-AFC, Ac-VEID-AFC, Ac-LETD-AFC, Ac-LEHD-AMC, Ac-IETD-AMC), the samples were excited at 400 nm (AFC substrates) or 350 nm (AMC substrates), and fluorescence emission was monitored at 505 nm (for AFC substrates) or 450 nm (for AMC substrates) for 60 seconds using a PTI fluorometer (Photon Technology International, Edison, NJ, USA). The steady-state parameters, K_M_ and k_cat_, were determined from plots of initial velocity *versus* substrate concentration.

Substrate phage display assays were performed as described (36, 37). Briefly, phage libraries consisting of caspase recognition sequences were bound to Ni-NTA resin. Enzyme (10-100 nM) was added to initiate the reaction, and samples were incubated between 3 and 20 hours. *E. coli* ER2738 cells were used to amplify the cleaved phage from previous rounds by infecting cells with the supernatant after enzyme incubation. The cells were grown for 4 hours, removed by centrifugation, and the supernatant was collected and used as the library for the following round of selection. Plaque counting was used to determine the endpoint of the experiment, when the number of phage bound to the resin was similar to the number of phage released during the treatment. The number of phage released during the reaction *versus* the control (without enzyme) was monitored to ensure progress in substrate selectivity.

### X-ray Crystallography

Protein structure predictions were performed using Swiss-Model (38) using human caspases-3, −6, and −7 as references (PDB ID 2J30, 3OD5, and 1F1J, respectively). For structure determination, the coral caspase proteins were dialyzed in a buffer of 10 mM Tris-HCl, pH 7.9, 100 mM NaCl, 1 mM DTT and concentrated to ∼7 mg/mL. The molar extinction coefficients for the proteins were determined by ProtParam under reduced conditions (Supplementary Table S3). Inhibitor, Ac-DEVD-CHO (reconstituted in DMSO), was added at a 5:1 (w/w) inhibitor/protein ratio, and DTT and NaN_3_ were added to final concentrations of 10 and 3 mM, respectively. Samples were incubated for 1 hour in the dark on ice. Hanging-drop vapor diffusion method was applied using 4 µL drops that contained equal volumes of protein and reservoir solutions using the PEG/ion 2 screen (Hampton Research). PaCasp7a protein crystalized in a solution of 0.1 M sodium malonate pH 5.0, 12% w/v polyethylene glycol (PEG) 3350, and conditions were optimized such that the best diffracting crystals of PaCasp7a were obtained at 18 °C in a solution of 0.1 M sodium malonate, pH 4.9–5.1, 15–17% PEG 3350 (w/v), 10 mm DTT, and 3 mm NaN_3_. Crystals for PaCasp7a appeared within 3 to 5 days and were briefly immersed in a cryogenic solution containing 20% PEG 4000, 80% reservoir solution. Crystals were stored in liquid nitrogen. We were unable to obtain diffraction quality crystals for the remaining coral caspases. Data sets were collected at 100 K at the SER-CAT synchrotron beamline (Advance Photon Source, Argonne National Laboratory, Argonne, IL). Each data set contained 180 frames at 1° rotation. The protein crystallized in the space group P 2_1_ 2_1_ 2_1_ and was phased with a previously published HsCasp3 structure (PDB entry 2J30). Data reduction and model refinements were done using HKL2000, COOT, and Phenix, and a summary of the data collection and refinement statistics is shown in Supplementary Table S4. Molecular dynamics simulations were performed for 50 ns with GROMACS 4.5 (39) using the Amber99 force field (40) and the TIP3P water model (41), as previously described (42).

### Data Deposition

The crystal structure for PaCasp7a has been deposited in the Protein Data Bank, www.wwpdb.org under PDB ID code: 6WI4.

## RESULTS

### Caspases in two coral species: Phylogenetic analysis and domain organization

We examined seven caspase genes from *O. faveolata*, based on sequences obtained from previous transcriptomic and genomic data (Supplementary Fig. S1 and Supplementary Table S1) (20). The caspases were named based on the E-value from BLAST as well as the sequence similarity to the human orthologs. Results from examining the sequence homology and domain organization suggest that three of the caspases are apoptotic initiators and four are apoptotic effectors in *O. faveolata* (Fig. 1A). The sequence identities of the seven caspases compared to most human caspases are low, only ∼35% (Table 1), so it is difficult to determine the nature of each coral caspase based solely on sequence comparisons with human orthologs. In addition, two caspases from *O. faveolata* contain an N-terminal CARD (caspase activation and recruitment domain) motif, similar to those in HsCasp2 and HsCasp9, and one caspase contains tandem DED (death effector domain) motifs, similar to that found in HsCasp8 (Fig. 1A). The remaining four proteins show domain organization similar to the human effector caspases, with short pro-domains (Fig. 1A).

**Table 1:** Protein sequence identity/similarity (%) with human caspases.

|  | *HsCasp3* | *HsCasp7* | *HsCasp6* | *HsCasp2* | *HsCasp8* | *HsCasp10* | *HsCasp9* |
| --- | --- | --- | --- | --- | --- | --- | --- |
| OfCasp7 | 37/54 | 38/52 | 32/49 | 28/43 | 34/50 | 33/53 | 34/50 |
| OfCasp3c | 36/58 | 35/56 | 35/52 | 32/48 | 37/53 | 37/54 | 33/49 |
| OfCasp3b | 35/60 | 32/57 | 33/52 | 32/49 | 37/55 | 33/53 | 34/50 |
| OfCasp3a | 47/69 | 45/65 | 38/54 | 29/48 | 39/54 | 39/55 | 28/46 |
| OfCasp2 | 37/53 | 41/55 | 35/46 | 33/52 | 34/52 | 35/52 | 32/48 |
| OfCasp8a | 39/56 | 39/53 | 34/48 | 35/53 | 32/58 | 30/49 | 34/51 |
| OfCasp8b | 33/56 | 31/50 | 31/49 | 32/52 | 35/51 | 34/52 | 32/46 |
| PaCasp3 | 36/58 | 37/59 | 36/56 | 33/49 | 37/54 | 34/54 | 35/50 |
| PaCasp7a | 43/65 | 44/60 | 36/53 | 28/46 | 37/53 | 37/53 | 28/44 |
| PaCasp7b | 38/54 | 37/52 | 34/49 | 29/46 | 31/48 | 30/50 | 33/48 |
| PaCasp2 | 39/53 | 38/52 | 33/48 | 33/50 | 35/53 | 35/52 | 32/49 |

**Figure 1:**
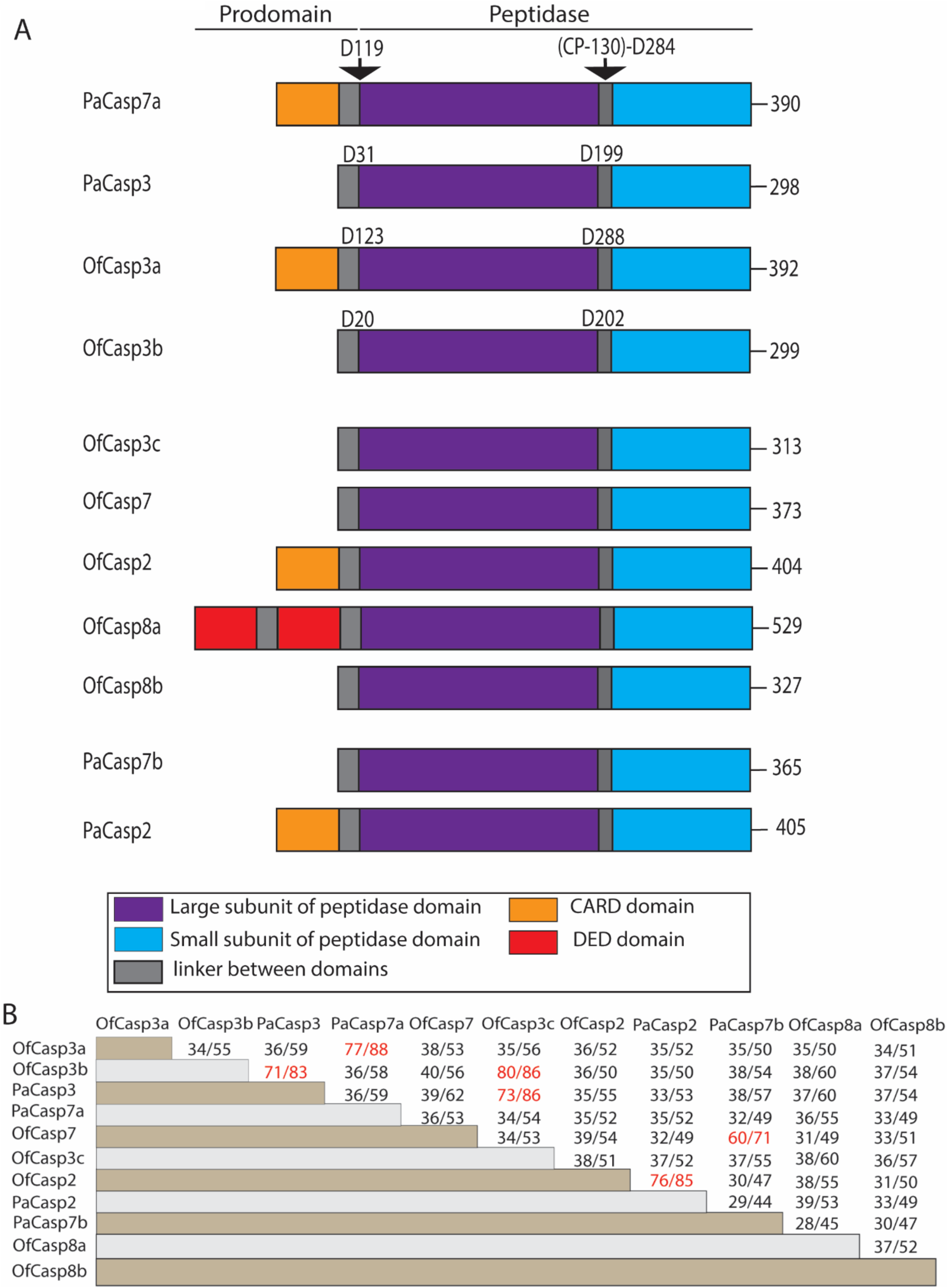
Domain organization and sequence comparison among caspases of *O. faveolata* and *P. astreoides*. (A) Domain organization of caspases in *O. faveolata* and *P. astreoides*. Processing site between large and small subunit, and after prodomain are noted in biochemically characterized caspase. (B) Protein sequence identity (%) and similarity (%) among coral caspases.

In the case of *P. astreoides*, four caspase sequences consisted of two initiator-like caspases (called PaCasp7a and PaCasp2) and two effector-like caspases (called PaCasp7b and PaCasp3) (Fig. 1A and Supplementary Fig. S1). Similar to the results for *O. faveolata*, the caspase sequences from *P. astreoides* also have only ∼35% identity with human caspases, regardless of comparisons to initiator or effector caspases (Table 1). The sequences from the two coral species displayed much higher identity to putative homologs in the other coral species. For example, PaCasp7a has a 77% sequence identity with OfCasp3a, whereas PaCasp3 has 71 and 73% sequence identity, respectively, with OfCasp3b and OfCasp3c. Likewise, PaCasp2 demonstrates 76% sequence identity with OfCasp2, and PaCasp7b shares 60% identity with OfCasp7 (Fig. 1B).

A phylogenetic analysis of cnidarian and vertebrate caspases demonstrated that cnidarian caspases cluster in separate groups (Fig. 2A). All of the short pro-domain caspases, including PaCasp3 and OfCasp3b, cluster together between vertebrate effector (caspases-3/7) and initiator (caspases-8/10) caspases. Interestingly, the comparative genomics and phylogenetic analyses suggest that short cnidarian caspases, that is, those lacking a CARD or DED, share a common ancestor with vertebrate effector caspases-3 and −7 and with initiator caspases-8 and −10 (Fig. 2A). Homologs of caspase-8 in coral share the same clade with vertebrate caspases-8 and −10, and the CARD-containing OfCasp2 and PaCasp2 clustered with vertebrate caspase-2. With the exceptions of OfCasp2 and PaCasp2, the other CARD-containing coral caspases cluster with OfCasp3a and PaCasp7a and segregate into a different clade, although they share a common ancestor with vertebrate caspases-2 and −9.

**Figure 2:**
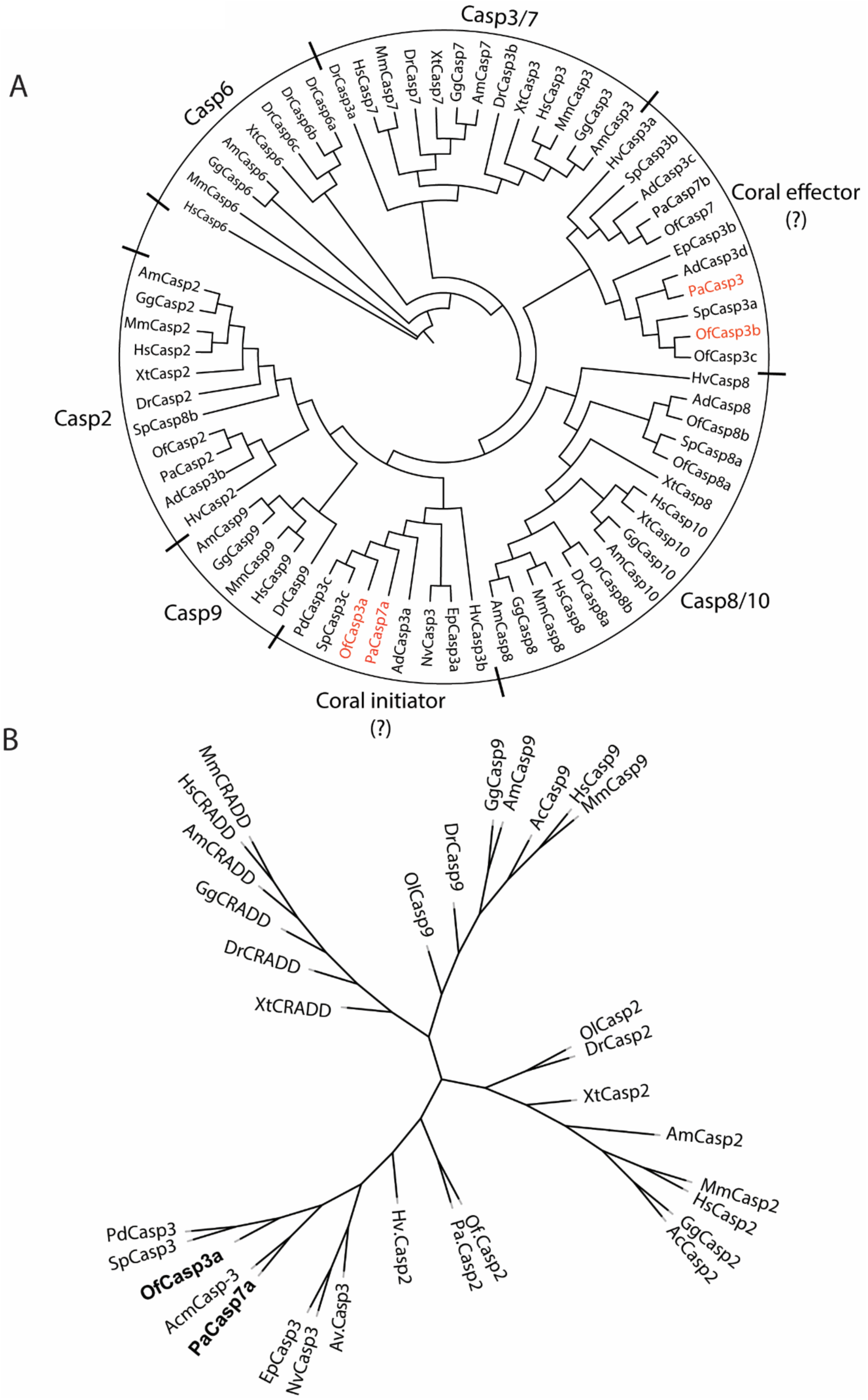
Coral caspase phylogenetic analysis. (A) Phylogenetic tree of cnidarian and vertebrate caspases. *Orbicella faveolata* (Of), *Porites astreoides* (Pa), *Pocillophora damicornis* (Pd), *Stylophora pistillata* (Sp), *Nematostella vectensis* (Nv), *Exaiptasia pallida* (Ep), *Hydra vulgaris* (Hv), *Acropora digitophora* (Ad), *Homo sapiens* (Hs), *Mus musculus* (Mm), *Gallus gallus* (Gg), *Alligator mississippiensis* (Am), *Xenopus laevis* (Xl), *Danio rerio* (Dr). Accession number of all used sequences are shown in Supplementary Tables S1 and S2. (B) Phylogenetic analysis of CARD domains of caspases and CRADDs (CASP2 and RIPK1 Domain containing Adaptor with Death Domain) between cnidarians and vertebrates.

We analyzed the CARD motifs of cnidarian caspases independently of the protease domains and compared them to the CARD motifs of vertebrate caspases-2 and −9 as well as that of CRADD (<u>c</u>aspase-2 and <u>R</u>IPK1 domain containing <u>a</u>daptor with <u>d</u>eath <u>d</u>omain) motifs, which recruits caspase-2 to the PIDDosome (43) (Fig. 2B). The CARD motifs of coral caspases-3 and - 7 cluster together but are more closely related to the CARD of caspase-2 than those of caspase-9 or CRADD. Based on this analysis, there appear to be many CARD-containing caspase-3-like proteins in cnidaria. At present, it is not clear why CARD-containing caspase-3-like proteins provide an advantage for coral development and/or symbiosis since the animals also contain initiator caspases that presumably activate the short pro-domain effector caspases. CARD-containing caspase-3-like proteins are rarely observed in vertebrate effector caspases. Fish-specific caspases have been found, such as the CARD-containing caspase-8 for example (44), but caspase-2 is, at present, the only characterized DxxDase with a CARD.

We chose two caspases from each species to characterize further, based on the sequence comparisons with human effector caspases-3, −6, or −7. In the case of *O. faveolata*, we chose two caspase-3-like proteins that showed 47% and 35% sequence identity, respectively, with HsCasp3, and we named the two proteins OfCasp3a and OfCasp3b, respectively (Fig. 1A and Table 1). Interestingly, despite predicted similarity to HsCasp3, OfCasp3a also has an N-terminal CARD motif. One caspase from *P. astreoides* demonstrated the highest sequence identity with HsCasp7 (44%) and was named PaCasp7a, even though it also contains a CARD motif (Fig. 1A and Table 1). The second protein from *P. astreoides* showed similar sequence identity to human caspases-3, −6, −7, and −8 (36-37%) (Fig. 1A and Table 1), but the protein does not have a DED motif like caspase-8 and the domain organization is more similar to that of caspase-3. Consequently, we named the protein PaCasp3. Overall, the low sequence identity between the vertebrate and invertebrate caspases show that the classification is somewhat arbitrary without further biochemical characterizations of the proteins. Together, the phylogenetic analysis shows that the caspases from *P. astreoides* and *O. faveolata* have relatively low sequence identity (∼40%) to mammalian caspases as well as other vertebrate families, but the proteins had much higher sequence identities to caspases from other cnidarian species, such as *Pocillopora damicornis, Stylophora pistillata*, and *Nematostella vectensis*.

An analysis of the coral caspase sequences shows that the proteins contain all of the conserved features that define a caspase. For example, each protein contains the catalytic dyad, histidine (CP-075) and cysteine (CP-117) (Fig. 3), where “CP” refers to the common position defined previously for caspases (28). The conserved sequence that contains the catalytic histidine (CP-115)-QACRG-(CP-119) is found in the four coral caspases, although PaCasp7a and OfCasp3a contain QACQG as in human caspase-8. One of the most highly variable regions, the intersubunit linker (IL) is the same length in OfCasp3b and PaCasp3 compared to that of HsCasp3, while those of PaCasp7a and OfCasp3a have 1 and 2 amino acids fewer than HsCasp3 respectively (Fig. 3).

**Figure 3:**
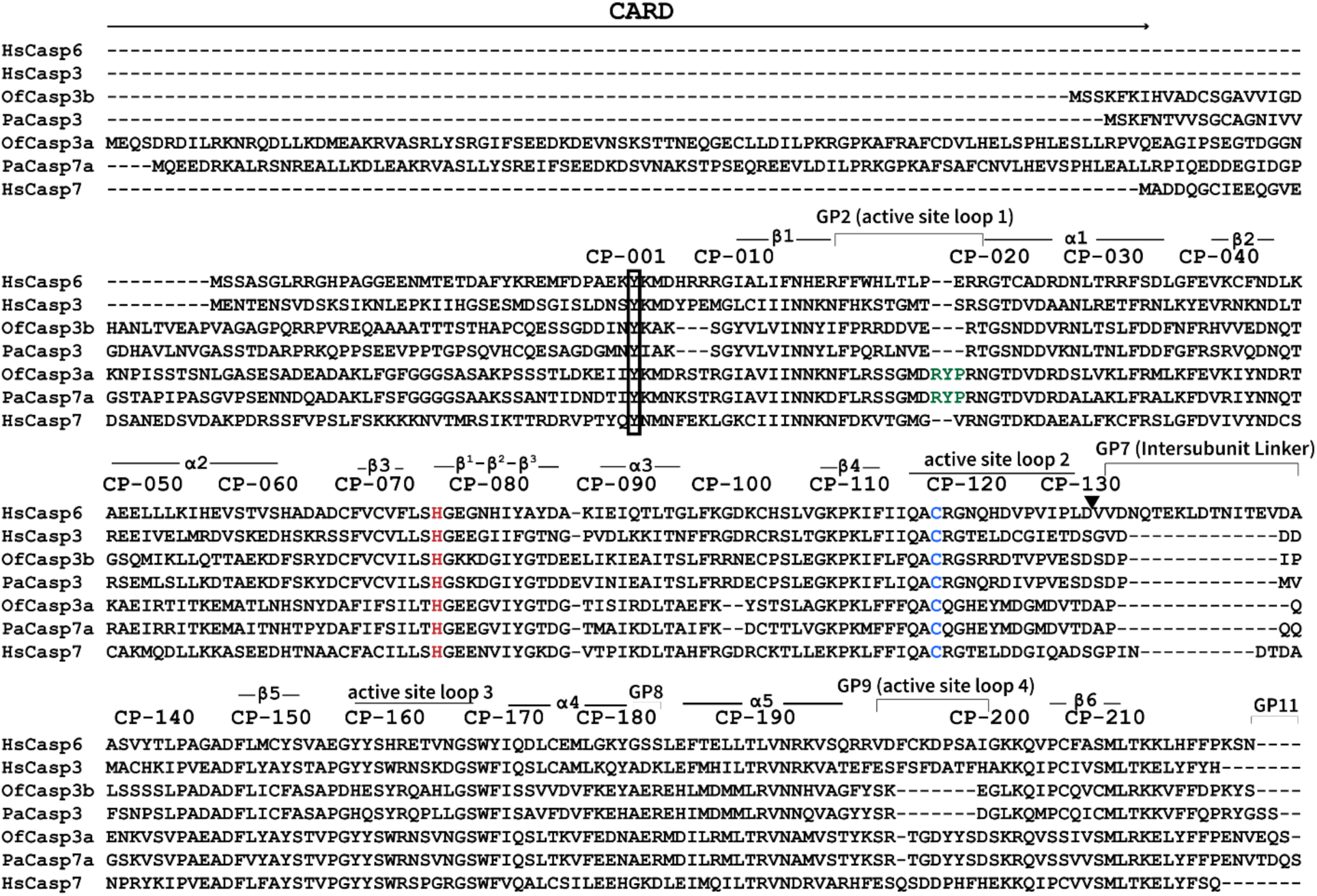
Two caspases each of *O. faveolata* and two of *P. astreoides* sequences aligned with human effector caspases. In the multiple sequence alignment (MSA), secondary structures (alpha helices, beta sheets, and loops) are indicated along with common position (CP) numbers among caspases. Gap positions, or sequences between common amino acid positions, are referred to as GP. Histidine (H) and Cysteine (C), which forms a catalytic dyad, are colored in red and blue respectively. “RYP” motif insertion in OfCasp3a and PaCasp7a are colored in green.

### Biochemical characterization of coral caspases

We examined the four coral caspases by size exclusion chromatography (SEC) since CARD-containing human caspases are monomers or mixtures of weak protomer-dimer (45). Because the IL of the procaspase monomer is cleaved during activation, the protomer is defined as a single unit that contains a large and small subunit and a single active site. Thus, the dimer consists of two protomers, or is more formally considered a dimer of heterodimers. The data show that the CARD containing coral caspases, PaCasp7a and OfCasp3a, elute in a single peak with MW of 42.6 and 44 kDa, respectively. The sizes are larger than that of a protomer but smaller than a dimer (Supplementary Fig. S2 and Supplementary Table S5) suggesting that the proteins form weak dimers similar to the human initiator caspases. In contrast, the short pro-domain containing caspases, PaCasp3 and OfCasp3b, are dimers similar to the human effector caspases, with MW of 64.5 and 69.2 kDa, respectively (Supplementary Fig. S3 and Supplementary Table S5).

We also determined the mass of the large and small subunits by mass spectrometry. Caspase zymogens are cleaved in the IL, and the N-terminal CARD or pro-domain is removed during activation (45). The proteins also auto-process during overexpression in *E. coli*. The MW of the large and small subunits of each caspase, determined by MS, are shown in Supplementary Table S5. When compared to the sequences for each protein (Fig. 3), the data show that OfCasp3a and PaCasp7a are cleaved in the intersubunit linker after (CP-127)-DVTD-(CP-130), whereas OfCasp3b and PaCasp3 are cleaved after (CP-127)-VESD-(CP-130). The actual amino acid positions, in addition to the common position number, are shown in Fig. 1A, and the cleavage sites are indicated by the arrow in Fig. 3. In addition, the first twenty or thirty-one amino acids, respectively, in the prodomains of OfCasp3b and PaCasp3 are removed following cleavage after VIGD (D^20^) (OfCasp3b) or SSTD (D^31^) (PaCasp3). The CARD motifs of OfCasp3a and of PaCasp7a are removed following cleavage after DEAD (D^123^) and DQAD (D^119^), respectively (Fig. 1A and Fig. 3). We note that there are potentially other cleavage sites in the CARD motifs, but in our assays the CARD motif was completely removed.

We characterized the substrate specificity for each of the four coral caspases using substrate-phage display assays, as described previously (37). In these assays, we utilize two substrate-phage libraries that determine the P5-P1’ substrate preferences, with either aspartate fixed at the P1 position (P5-xxxxDx-P1’) or random (called 6x), and the results were the same for both libraries. The data show that PaCasp7a and OfCasp3a have Group II specificity, with a preference for aspartate in the P4 position (DxxDase) (Fig. 4A and Fig. 4B). In contrast, PaCasp3 and OfCasp3b prefer valine in the P4 position (VxxDase) (Figs 4C and 4D), which is defined as Group III specificity like HsCasp6.

**Figure 4:**
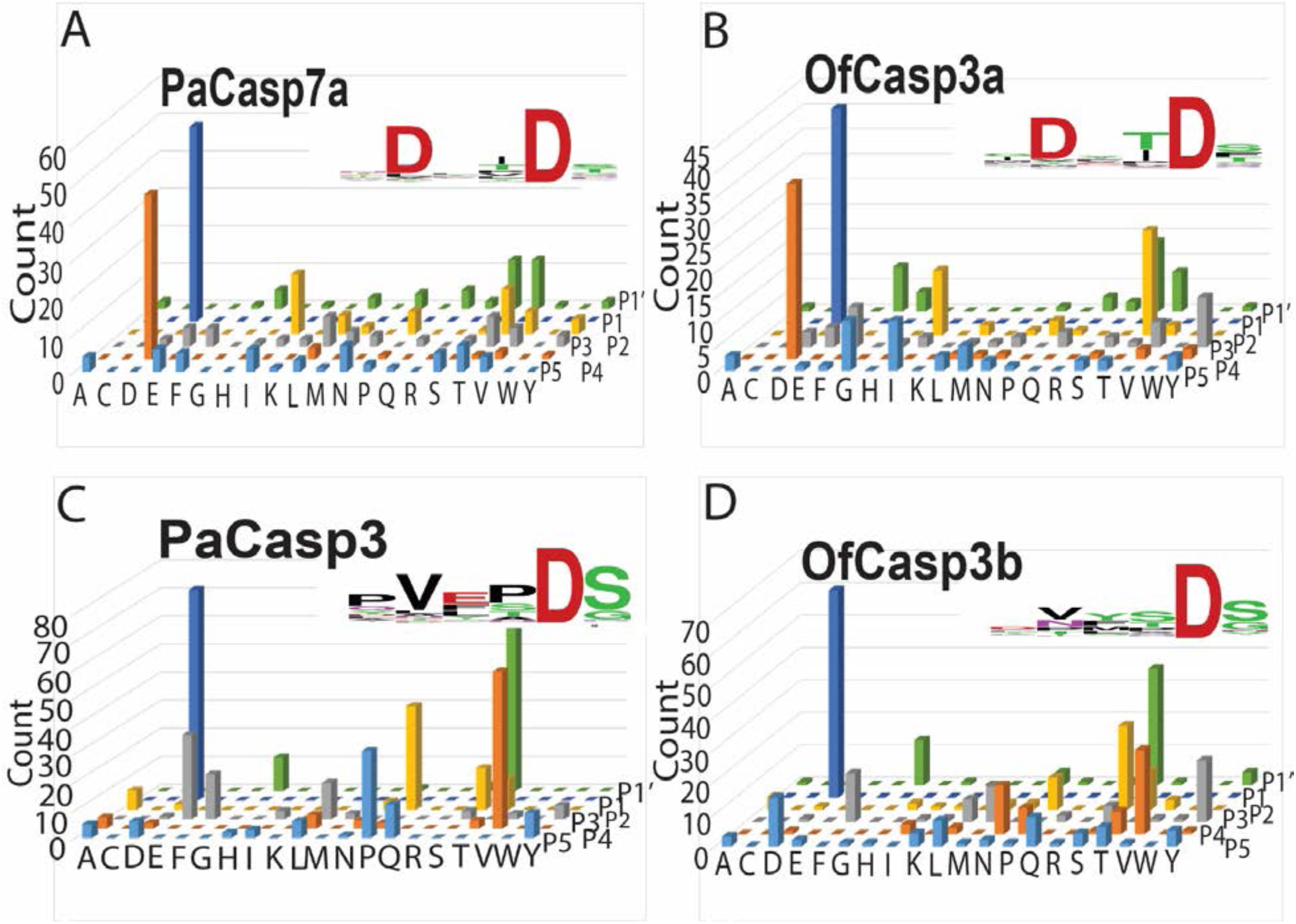
Substrate preference determined by substrate-phage display. Amino acid preferences shown for substrate positions P5-P4-P3-P2-P1-P1’ for PaCasp7a (A), OfCasp3a (B), PaCasp3 (C) and OfCasp3b (D). Values of Y-axes indicate number of phage sequences containing the specified amino acid. Web logos are also shown in inset of respective graph for same results.

The activities of PaCasp7a and of OfCasp3a were also examined using DEVD-AFC and VEID-AFC substrates. In all cases, however, the activity against the tetrapeptide substrates was very low due to K_M_ values >500 μM, so we could not reliably determine the steady-state catalytic parameters k_cat_ or K_M_ from the small peptide activity assays. In caspases, the K_M_ is thought to correlate with substrate binding (K_D_), so the high K_M_ suggests poor binding of the small peptide.

Because of the low activity in small peptide assays, we tested the coral caspases for their ability to hydrolyze full-length (FL) human procaspases-3 and −6, which were made catalytically inactive due to mutation of the catalytic cysteine to serine (26). Thus, the proteins are incapable of undergoing self-proteolysis. As shown in Fig. 3, HsCasp3 is cleaved once in the intersubunit linker at CP-130 (IETD), while HsCasp6 contains two cleavage sites at CP-130 (DVVD) and at GP7-D17 (TEVD). Each procaspase substrate was incubated separately with an active coral caspase, and the reaction was monitored over eight hours. Aliquots were removed and analyzed by SDS-PAGE (Fig. 5A). The results show that procaspase-3 was cleaved by PaCasp3 and by OfCasp3b, with little to no cleavage by PaCasp7a or by OfCasp3a. In contrast, procaspase-6 was cleaved by PaCasp7a and by OfCasp3a, but there was little to no cleavage by PaCasp3 or by OfCasp3b. Together, the data corroborate our results from substrate-phage display (Fig. 4) that identify PaCasp3 and OfCasp3b as VxxDases and PaCasp7a and OfCasp3a as DxxDases, respectively.

**Figure 5:**
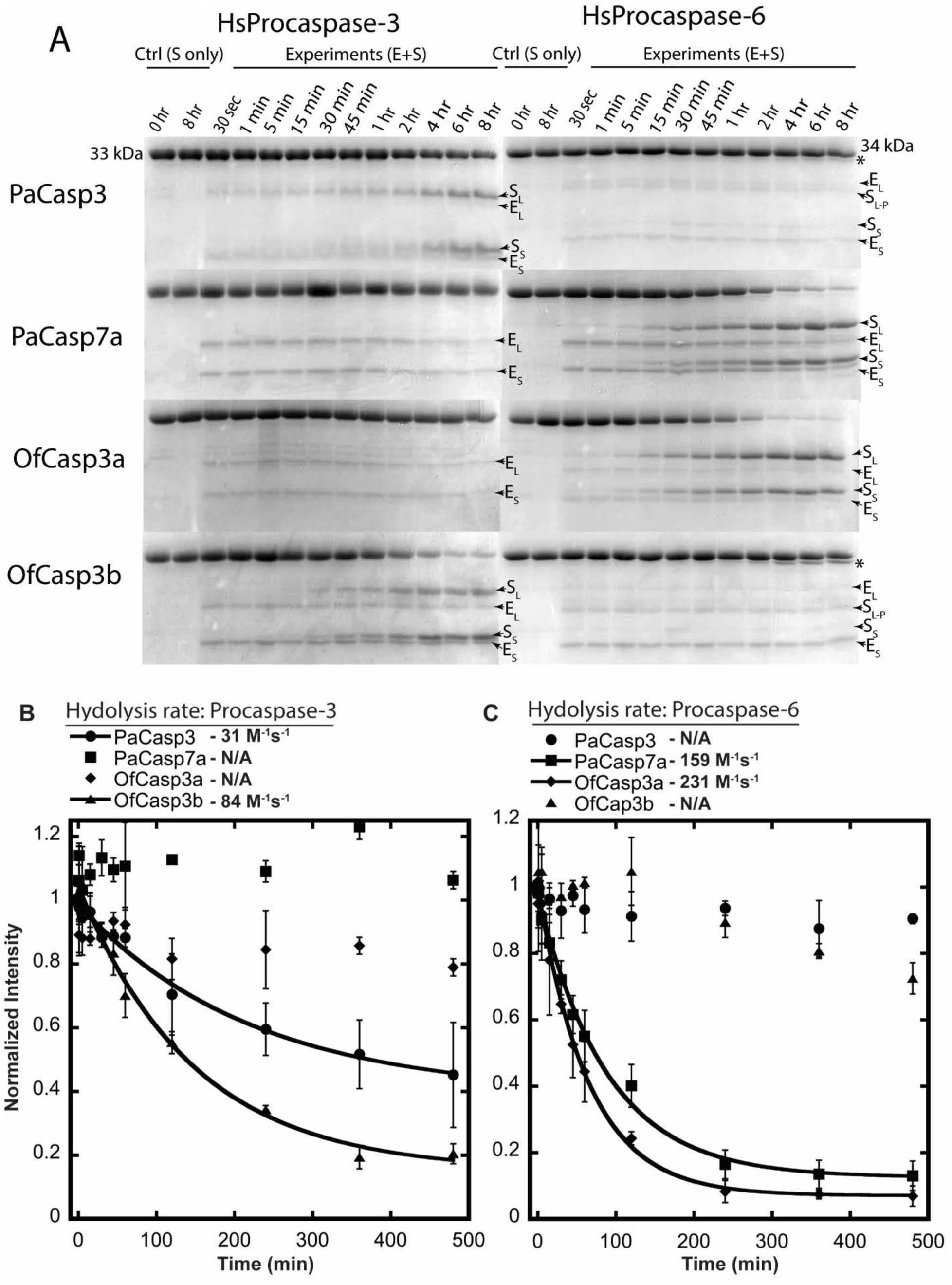
Cleavage kinetics of coral caspases using human procaspases-3 and −6 as a substrate. (A) Cleavage of full-length inactive HsCasp3 and HsCasp6 by coral caspases over time course. All cleaved products are labeled along with enzyme itself. (S_L_- large subunit of substrate, S_S_- small subunit of substrate, E_L_- large subunit of enzyme, E_S_- small subunit of enzyme, S_L-P_- large subunit with prodomain cleaved, S- substrate and E+S- enzyme and substrate). Bands with “*” indicate only prodomains were removed from full-length substrate. (B, C) Quantification of procaspase bands relative to the control (substrate without enzyme after 8 hours incubation). Data were fit to a single exponential decay to calculate CF_50_ used to calculate hydrolysis rate of coral caspases (solid line). Procaspase-3 (B), Procaspase-6 (C). Error bars represent standard deviation form three different experiments.

As described previously (34), we quantified the rate of hydrolysis of the two procaspase substrates by assessing the disappearance of the full-length procaspases-3 and −6, both ∼32 kDa in size, and the appearance of the large (∼20 kDa) and small (∼10 kDa) subunits over the time course of the assay (Fig. 5B and Fig. 5C). The data were fit to a single exponential decay to approximate k_cat_/K_M_. The results show that PaCasp3 and OfCasp3b cleave procaspase-3 with hydrolysis rates of 31 M^-1^s^-1^ and 84 M^-1^s^-1^, respectively (Fig. 5B), while PaCasp7a and OfCasp3a cleaved procaspase-6 with hydrolysis rates of 159 M^-1^s^-1^ and 231 M^-1^s^-1^, respectively (Fig. 5C). We note that, although not quantified, both PaCasp3 and OfCasp3b cleave the procaspase-6 pro-peptide (TETD) at a much slower rate than that observed for cleavage of the intersubunit linker of procaspase-3 (IETD). Together, the biochemical data show that the coral caspases are weak enzymes, at least in the *in vitro* assays, with k_cat_/K_M_ values ∼10^2^ M^-1^s^-1^.

### Crystal structure of PaCasp7a

We attempted to crystalize all four of the coral caspases, and we were successful in obtaining diffraction quality crystals of PaCasp7a with an inhibitor (DEVD-CHO) bound in the active site. The crystals diffracted in the P2_1_2_1_2_1_ space group, and we determined the structure to high resolution at 1.57 Å (Supplementary Table S4). The data show that the PaCasp7a is very similar to human caspases, with an RMSD of <1 Å compared to HsCasp3 (Fig. 6A). In the active site, the carboxylate group of the P4 aspartate hydrogen bonds to Asn^315^ (CP-162) on active site loop 3 (L3), the backbone amide of Arg^356^ (GP9-02) on L4, and through-water hydrogen bonds to Trp^321^ (CP-168) (on L3) as well as the backbone carbonyl of Arg^356^ (GP9-02) (on L4) (Fig. 6B). In general, the active site provides hydrophilic binding pockets for the P3 glutamate and P4 aspartate of the substrate, and a more hydrophobic binding pocket for the P2 valine side-chain (Fig. 6C), similarly to that of HsCasp3.

**Figure 6.**
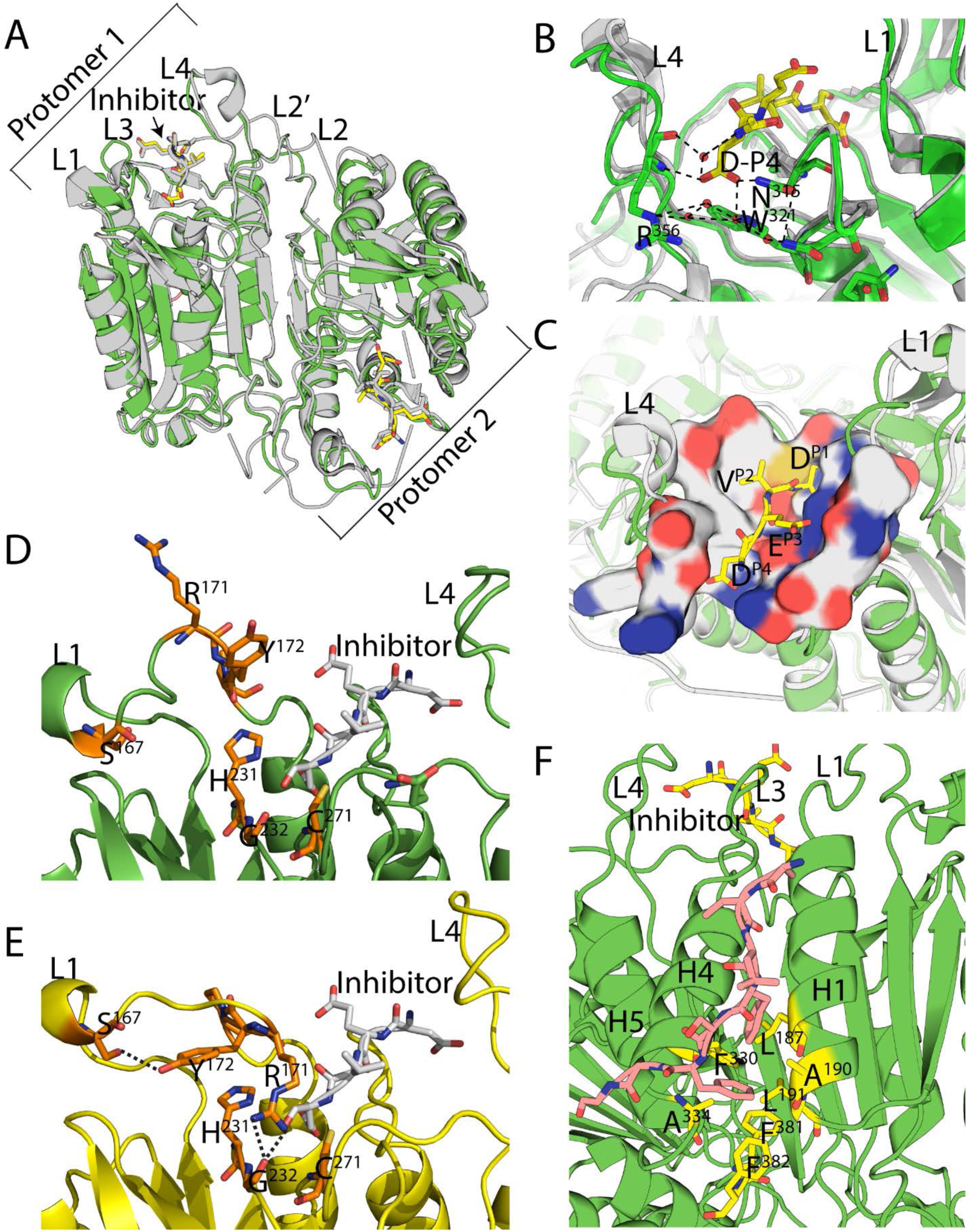
Structure of PaCasp7a. (A) Comparison of PaCasp7a (green) aligned with HsCasp3 (grey) (PDB ID: 2J30). (B) PaCasp7a active site bound with inhibitor DEVD-CHO. Dashed lines show hydrogen bonding network to the P4 aspartate. (C) Surface map of active site residues in PaCasp7 within 5 Å of the inhibitor (yellow sticks). Neutral charges are grey, negative charges are red, and positive charges are blue. (D) **“**Out” orientation, of “RYP” residues in loop 1 in crystal structure of PaCasp7a. (E) “In” orientation, of “RYP” residues in loop 1 in predicted model of PaCasp7a. Dashed lines show the hydrogen bonds formed by “R” and “Y” in “In” Orientation. (F) N-terminal peptide (orange) bound in hydrophobic pocket between helices 1 and 4. PaCasp7a residues that form the pocket are shown in yellow: L^187^ (CP-031), A^190^ (CP-034), L^191^ (CP-035), F^330^ (CP-177), A^334^ (CP-181) as well as F^381^ (CP-217) and F^382^ (CP-218) at the C-terminus.

Both PaCasp7a and OfCasp3a contain a two-residue insertion in loop 1 (L1) of the active site (Fig. 3). The structure of PaCasp7a with inhibitor bound shows that the insertion extends the loop compared to HsCasp3 and results in an “RYP” motif in L1 (Fig. 3) near the catalytic histidine (Fig. 6D). Models of the PaCasp7a active site suggest that rotation of the loop results in intercalation of the Arg^171^ (GP2-17) between the catalytic His^231^ (CP-075) and Cys^271^ (CP-117) (Fig. 6E). In this orientation, the arginine side-chain hydrogen bonds with the carbonyl of Gly^232^ (CP-076) on active site loop 3 (L3) and clashes with the P1 and P2 positions of substrate. In addition, the tyrosine of the RYP motif forms a new hydrogen bond with the side-chain of Ser^167^ (GP2-05) in L1 (Fig. 6E). Altogether, the models suggest that in the absence of substrate, rotation in L1 may stabilize an inactive conformation of the enzyme. We note, however, that MD simulations (50 ns) of the structural models show that the region of L1 that contains the RYP motif is very mobile, so if the RYP motif is indeed autoinhibitory, then the “RYP-In” conformation appears to be transient (Supplementary Fig. S3).

The structure of PaCasp7a also reveals a peptide bound on the protein surface near *α*-helices 1 and 4. The structure shows that amino acids in the N-terminus of PaCasp7a (N’-AKLFSFGG-C’) (N’-PD-A025 to PD-G018-C’ in the common position numbering) comprise the peptide, where the two phenylalanine side-chains bind in a hydrophobic pocket between the helices 1 and 4 (Fig. 6F). The binding pocket on the protein is formed by five hydrophobic residues on the two helices (L^187^ (CP-031), A^190^ (CP-034), L^191^ (CP-035), F^330^ (CP-177), A^334^ (CP-181)) as well as F^381^ and F^382^ (CP-217 and CP-218) at the C-terminus (Fig. 6F). The peptide also forms several hydrogen bonds with charged groups on the protein surface. We do not observe electron density for amino acids G^128^ (PD-017) - N^141^ (PD-004) (Fig. 3), but extensive interactions downstream of D^142^ (PD-003) result in an ordered structure that moves into the core of the protein. The fourteen disordered residues would provide ample distance to connect the peptide with the protease domain, and the data suggest that the intervening amino acids may hinder dimerization since they would be anchored near the dimer interface when the peptide is bound on the protein surface. The N-terminal end of the peptide is immediately downstream of the DQAD cleavage site that removes the CARD motif (Fig. 3), suggesting that the binding pocket on the protein surface may be used to position the N-terminal linker (between the CARD and protease domains) in the active site.

We searched the caspase structures in the protein data bank and found four examples of an N-terminal peptide bound between helices 1 and 4 – human caspases-1 (PDB ID 2H48) and −2 (PDB ID 3R7M), Dronc of *Drosophila melanogaster* (PDB ID 2FP3) and CED-3 of *Caenorhabditis elegans* (PDB ID 4M9R) (46–49) (Supplementary Figs. S4A – S4D). Interestingly, the region of the peptide that is disordered in PaCasp7a (G^128^ PD-017 - N^141^ PD-004) forms a short *α*-helix in caspases-1 and −2 and in CED-3 (Supplementary Figs. S4A – S4B and S4D). The short helix does not make contacts across the dimer interface but rather makes extensive intra-protomer contacts with the C-terminus of the protein. In the case of DRONC, the intervening peptide forms an extended structure that extends beyond the dimer interface and would clash with the second protomer of the dimer (Supplementary Fig. S4C). In all cases, the binding pocket between helices 1 and 4 is hydrophobic, and the peptide binds through insertion of one or more hydrophobic amino acids into the binding pocket as well as hydrogen bonds between side-chains on the protein surface and backbone atoms of the peptide. Therefore, the structures show a common theme in which the N-terminal peptide downstream of the pro-domain cleavage site binds to a hydrophobic pocket on the protein surface. The interactions likely stabilize the peptide in the binding pocket for cleavage.

Finally, we observed a similar hydrophobic pocket in human effector caspases (HsCasp3, PDB: 2J30; HsCasp6, PDB: 3S70; HsCasp7, PDB: 1F1J) (50–52) (Supplementary Figs. S4E – S4G). There is no evidence, however, from biochemical or structural data, that the N-terminal peptide of the short pro-domain caspases bind in the hydrophobic pocket. A comparison of the N-terminal sequences (Fig. 3) shows significant divergence in the peptide of human effector caspases, so although the binding pocket is similar to that of PaCasp7a, the binding interactions with the peptide sequences are not similar. In HsCasp3 and HsCasp6, for example, the cleavage site is downstream of the putative binding sequence, so the entire peptide is removed from the N-terminus. Interestingly, in HsCasp7 the cleavage site is upstream of the binding region, but the sequence evolved into a tetra-lysine motif that has been shown to be an exosite for substrate selection in caspase-7 (53).

## DISCUSSION

Coral reefs are facing a significant decline due to increasing local and global stressors from disease, climate change, and pollution (54). While coral possess a robust innate immune system, the cellular responses to disease have not been elucidated, beyond generalized major response categories. In addition, elevated ocean temperatures have emerged as key threats to the long-term survival of coral reefs and are leading to a collapse of the coral-algal symbiosis (21, 54). The algal symbionts are responsible for about 90% of the coral metabolic needs, so the mortality rate of bleached coral is high (55, 56). Despite the environmental consequences posed by coral disease and coral bleaching, and resultant changes to reef communities, the molecular physiology behind coral immune responses is not well understood (57–60). A better understanding of coral stress and immune mechanisms, including cell signaling and biochemical response mechanisms, will improve our understanding of coral declines. The two coral species described here, *Orbicella faveolata* and *Porites astreoides* are both reef-building coral, but the two species lie on opposite ends of the disease response spectrum. Where *O. faveolata* is sensitive to disease and activates apoptotic responses to stress, *P. astreoides* is resistant to disease and activates autophagic responses to stress. Thus, the two stony coral represent intriguing systems to characterize the biochemical responses to stress (18).

We show here that the coral caspases have relatively low sequence identity to human caspases, so the designation of the caspase function is somewhat arbitrary without further biochemical characterization. The data suggest that PaCasp7a and OfCasp3a may function similarly to caspase-2 since they exhibit DxxDase activity and contain an N-terminal CARD motif. In contrast, PaCasp3 and OfCasp3b share characteristics with effector caspase-6, with a short prodomain and VxxDase activity. Moreover, a phylogenetic analysis showed that OfCasp3a and PaCasp7a are close to vertebrate initiator caspases, whereas PaCasp3 and OfCasp3b are closer to effector caspases. Although the caspases exhibited low activity against peptide substrates, we were able to confirm the selection through cleavage assays of protein substrates. The results showed that the DxxDases (PaCasp7a and OfCasp3a) processed procaspase-6, which has a DVVD cleavage sequence in the intersubunit linker, but not procaspase-3, which contains a more hydrophobic recognition sequence recognized by caspase-8 (IETD). The opposite was true for PaCasp3 and OfCasp3b. In those cases, the enzymes processed procaspase-3 but not procaspase-6.

Taken together, the biochemical data show that the two short prodomain caspases (OfCasp3b and PaCasp3) are most likely not the main executioner caspases in coral. Rather, the enzymes may function similarly to HsCasp6 during cell development. In addition, the two proteins are not directly activated by the CARD-containing caspases, PaCasp7a or OfCasp3a. At present, it is not clear if the DxxDase activity of the two enzymes functions as the primary executioner of apoptosis or if the proteins activate the as-yet-unidentified executioner caspase. Based on the caspases identified in coral, we suggest a model in which the PaCasp7a and OfCasp3a enzymes are activated on PIDDosome-like complexes, similar to HsCasp2 (Figure 7). Either the DxxDase activity is utilized to kill cells, like CED-3 in *C. elegans*, or the activated enzymes cleave the executioner. In the latter case, the PaCasp3 and OfCasp3b proteins would be indirectly activated by PaCasp7a and OfCasp3a, respectively, through the undefined executioner caspase. Alternatively, the OfCasp3a and PaCasp7a may be activated on apoptosome complexes, and the DxxDase activities could be used to activate downstream caspases or to execute apoptosis. The latter suggestion is consistent with the presence of coral caspase-2-like proteins that also contain CARD motifs (Fig. 1A). Caspase-8-like proteins containing DED motifs would also activate the executioner (and PaCasp3 or OfCasp3b) indirectly. The putative caspase-8 and caspase-3-executioner proteins have been identified, but not yet characterized, in coral (25, 61–63). In addition, PIDDosome complex proteins are also present in coral (64). Together, the data suggest that coral are responsive to death ligands as well as metabolic changes in the cell. Further characterization of the apoptotic components will determine signaling responses leading to activation of the caspases, in particular whether OfCasp3a and PaCasp7a are activated on PIDDosome or apoptosome complexes.

**Figure 7.**
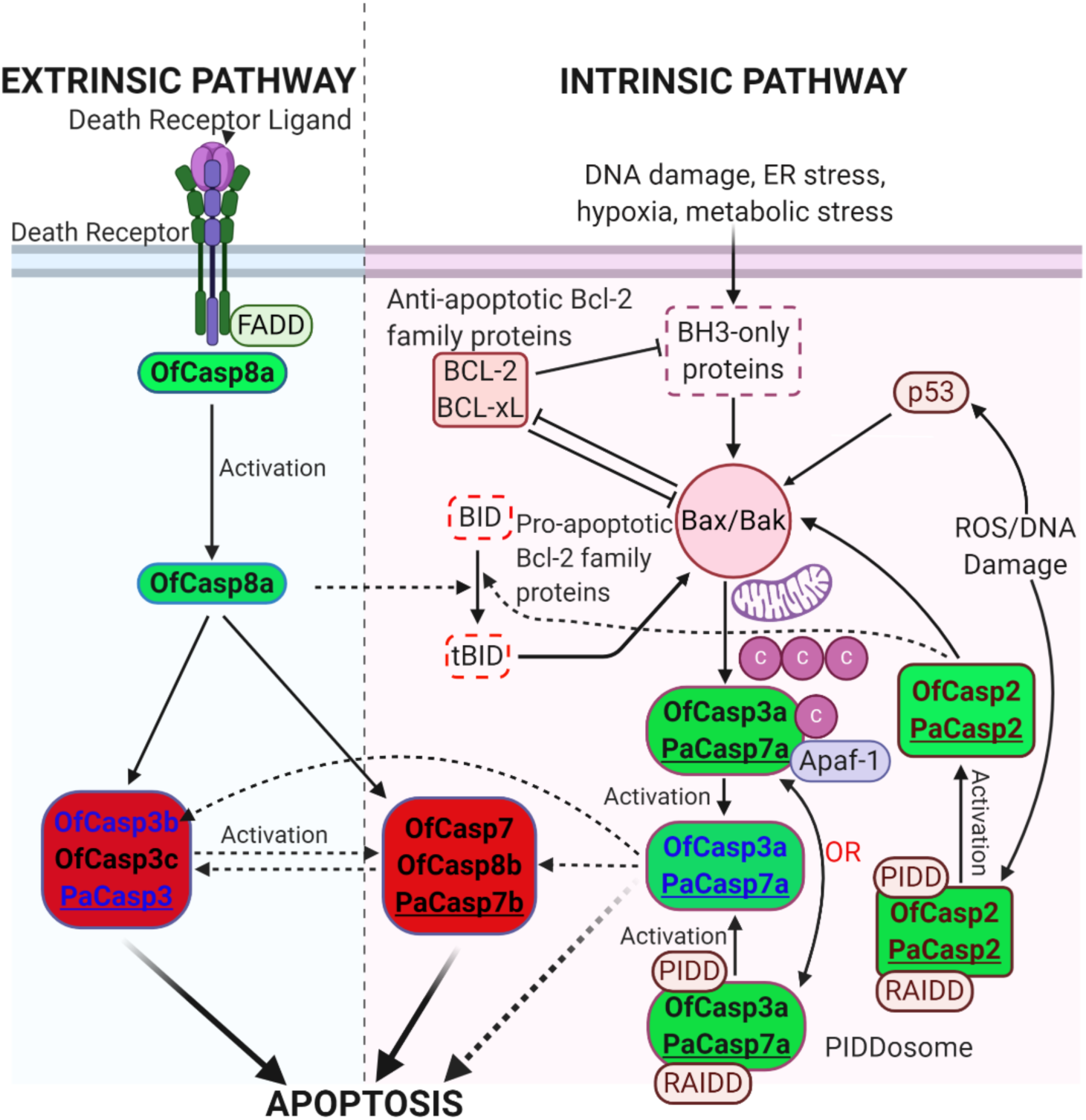
Proposed apoptotic pathways in coral compared to the apoptotic pathways in humans. All components in the pathways have homologs in *O. faveolata* and *P. astreoides* with the exception of BID (dotted box). A list of homologs is shown in Supplementary Table S6. Dotted lines indicate that links have not yet been shown experimentally. Caspases in green background are initiators and those in red background are effectors. Pa refers to *P. astreoides* and Of refers to *O. faveolata.* The four caspases characterized here are shown in blue.

Finally, the data shown here for PaCasp7a, as well as previous structural data – human caspases-1 (PDB ID 2H48) and −2 (PDB ID 3R7M), Dronc of *Drosophila melanogaster* (PDB ID 2FP3) and CED-3 of *Caenorhabditis elegans* (PDB ID 4M9R), and human effector caspases (HsCasp3, PDB: 2J30; HsCasp6, PDB: 3S70; HsCasp7, PDB: 1F1J) (46–52) (Supplementary Figs. S4A – S4G) – identify a hydrophobic pocket on the protein surface between helices 1 and 4 in which a peptide sequence C-terminal to the processing sequence binds. The binding of the peptide may increase activity through improved binding of the recognition sequence in the active site and help position the linker near the CARD-motif for cleavage. Alternatively, the binding of the peptide to the pocket may affect substrate selection by demonstrating a preference for substrates with both a P1-P4 cleavage sequence and the downstream sequence that binds to the pocket. We also showed that the hydrophobic pocket is conserved in a wide range of species, with similar size and properties. The short pro-domain caspases appear to have retained the binding pocket on the protease domain, but the N-terminal peptide sequence diverged, suggesting that effector caspases may utilize the binding pocket as an exosite for substrate selection. In this case, for example, substrates with sequences that bind in the pocket, and are downstream of the cleavage site, may exhibit better binding compared to substrates that contain only the P1-P4 recognition sequences.

## Conclusions

Coral have complex apoptotic signaling cascades, similar to those of vertebrates. We have identified OfCasp3a and PaCasp7a as initiator caspases that appear to function similarly to HsCasp2, indicating that coral are responsive to metabolic changes in the cell. In addition, both *P. astreoides* and *O. faveolata* contain VxxDases similar to HsCasp6. Our data show that the enzymes from both species have similar biochemical properties and are activated by similar mechanisms. Together, the data show that the role of the caspase cascade in disease resistance of *P. astreoides* or in disease sensitivity of *O. faveolata* may derive from differences in response mechanisms in the death receptor or the PIDDosome activation platforms, that is, signaling events upstream of the caspase cascade. Since the caspases in the two species exhibit similar biochemical properties and activation mechanisms, our data suggest that differences in the receptor-mediated activation of caspases as well as cross-talk between the autophagic and apoptotic pathways in the two coral species lead to the different physiological responses.

## ACKNOWLEDGEMENT

This work was supported by a grant from the National Institutes of Health [grant number GM127654 (to A.C.C)] and by funds from UT Arlington [Office of the Vice President for Research (to A.C.C)]. Use of the Advanced Photon Source was supported by the U.S. Department of Energy, Office of Science, Office of Basic Energy Sciences, under contract number W-31-109-ENG-38.

