## Supplemental Figures Tables for "Caspases from Scleractinian Coral Show Unique Regulatory Features"

**Supplementary Table S1:** Caspases from O. faveolata and P. astreoides with their assigned name based on sequence and domain similarity with human caspases, their respective accession number, and domains in their structure.

| Given Name | Accession number | Domains |
|---|---|---|
| ***Orbicella faveolata*** | | |
| OfCasp3a | XP_020613409.1 | CARD, Peptidase_C14 |
| OfCas3b | XP_020630525.1 | Peptidase_C14 |
| OfCasp7 | XP_020613679.1 | Peptidase_C14 |
| OfCasp3c | XP_020630550.1 | Peptidase_C14 |
| OfCasp2 | XP_020630531.1 | CARD, Peptidase_C14 |
| OfCasp8a | XP_020620405.1 | DED, DED, Peptidase_C14 |
| OfCasp8b | XP_020629413.1 | Peptidase_C14 |
| ***Porites astreoides*** | | |
| PaCasp-3 | comp-76580 | Peptidase_C14 |
| PaCasp-7a | comp-74978 | CARD, Peptidase_C14 |
| PaCasp2 | comp-74936 | CARD, Peptidase_C14 |
| PaCas7b | comp-77018 | Peptidase_C14 |

**Supplementary Table S2:** Caspases from invertebrates and vertebrates used in the phylogenetic analysis and their respective accession number.

| Caspases name | Accession number |
|---|---|
| *Acropora digitifera* | |
| AdCasp3a | XP_015775441.1 |
| AdCasp3b | XP_015766400.1 |
| AdCasp3c | XP_015762449.1 |
| AdCasp8 | XP_015761120.1 |
| AdCasp3d | XP_015753767.1 |
| *Alligator mississippiensis* | |
| AmCasp6 | XP_019355646.1 |
| AmCasp8 | XP_006272599.1 |
| AmCasp10 | XP_014449789.1 |
| AmCasp7 | XP_014450146.1 |
| AmCasp2 | XP_014464708.1 |
| AmCasp3 | XP_019336883.1 |
| AmCasp9 | XP_019355521.1 |
| *Danio rerio* | |
| DrCasp2 | NP_001036160.1 |
| DrCasp3a | XP_001338890.2 |
| DrCasp3b | XP_005173133.1 |
| DrCasp6a | XP_005164109.1 |
| DrCasp6b | XP_017210076.1 |
| DrCasp6c | NP_001018333.1 |
| DrCasp7 | XP_005156389.1 |
| DrCasp8a | NP_571585.2 |
| DrCasp8b | NP_001092089.1 |
| DrCasp9 | NP_001007405.2 |

| | |
|---|---|
| *Exaiptasia pallida* | |
| EpCasp3a | XP_020893866.1 |
| EpCasp3b | XP_020905061.1 |
| *Gallus gallus* | |
| GgCasp2 | NP_001161173.1 |
| GgCasp3 | NP_990056.1 |
| GgCasp6 | NP_990057.1 |
| GgCasp7 | XP_421764.3 |
| GgCasp8 | NP_989923.1 |
| GgCasp9 | XP_424580.5 |
| GgCasp10 | XP_421936.4 |
| *Homo sapiens* | |
| HsCasp2 | NP_116764.2 |
| HsCasp3 | NP_004337.2 |
| HsCasp6 | NP_001217.2 |
| HsCasp7 | NP_001253985.1 |
| HsCasp8 | NP_001219.2 |
| HsCasp9 | NP_001220.2 |
| HsCasp10 | NP_116759.2 |
| *Hydra vulgaris* | |
| HvCasp2 | NP_001274285.1 |
| HvCasp3a | XP_012557085.1 |
| HvCasp3b | XP_002159783.3 |
| HvCasp7 | XP_012561656.1 |
| HvCasp8 | XP_012562456.1 |
| *Mus musculus* | |
| MmCasp2 | NP_031636.1 |
| MmCasp3 | NP_001271338.1 |
| MmCasp6 | NP_033941.3 |
| MmCasp7 | XP_006526679.1 |
| MmCasp8 | NP_001264855.1 |
| MmCasp9 | NP_056548. |
| *Nematostella vectensis* | |
| Nv.Casp3 | XP_001633895.1 |
| Pocillophora damicornis | |
| Pd.Casp3 | XP_027037576.1 |
| *Stylophora pistillata* | |
| SpCasp3a | XP_022784432.1 |
| SpCasp3b | XP_022808070.1 |
| Sp.Casp3c | PFX33553.1 |
| SpCasp8a | XP_022796790.1 |
| SpCasp8b | XP_022789601.1 |
| *Xenopus laevis* | |
| XtCasp2 | XP_012809163.1 |
| XtCasp3 | NP_001120900.1 |
| XtCasp6 | NP_001011068.1 |
| XtCasp7 | NP_001016299.1 |
| XtCasp8 | XP_017953067.1 |
| XtCasp10 | NP_001015715.2 |

**Supplementary Table S3:** Characteristics of coral caspases.

| Composition (1) | Protein | | | | | | |
|---|---|---|---|---|---|---|---|
| | OfCasp3a | OfCasp3b | PaCasp3 | PaCasp7a | HsCasp3 | HsCasp6 | HsCasp7 |
| Total Number Amino Acids | 392 | 299 | 298 | 390 | 277 | 293 | 303 |
| Ala (A) | 24 (6.1%) | 22 (7.4%) | 16 (5.4%) | 31(7.9%) | 12 (4.3%) | 19 (6.5%) | 18 (5.9%) |
| Arg (R) | 26 (6.6%) | 18 (6.0%) | 16 (5.4%) | 25 (6.4%) | 14 (5.1%) | 17 (5.8%) | 15 (5.0%) |
| Asn (N) | 19 (4.8%) | 11 (3.7%) | 16 (5.4%) | 19 (4.9%) | 15 (5.4%) | 11 (3.8%) | 14 (4.6%) |
| Asp (D) | 29 (7.4%) | 24 (8.0%) | 23 (7.7%) | 30 (7.7%) | 20 (7.2%) | 20 (6.8%) | 27 (8.9%) |
| Cys (C) | 3 (0.8%) | 9 (3.0%) | 9 (3.0%) | 3 (0.8%) | 8 (2.9%) | 10 (3.4%) | 11 (3.6%) |
| Gln (Q) | 10 (2.6%) | 10 (3.3%) | 14 (4.7%) | 12 (3.1%) | 4 (1.4%) | 7 (2.4%) | 11 (3.6%) |
| Glu (E) | 28 (7.1%) | 17 (5.7%) | 14 (4.7%) | 26 (6.7%) | 20 (7.2%) | 20 (6.8%) | 19 (6.3%) |
| Gly (G) | 25 (6.4%) | 16 (5.4%) | 20 (6.7%) | 23 (5.9%) | 16 (5.8%) | 19 (6.5%) | 18 (5.9%) |
| His (H) | 5 (1.3%) | 9 (3.0%) | 6 (2.0%) | 5 (1.3%) | 8 (2.9%) | 12 (4.1%) | 7 (2.3%) |
| Ile (I) | 21 (5.4%) | 16 (5.4%) | 15 (5.0%) | 21 (5.4%) | 19 (6.9%) | 13 (4.4%) | 17 (5.6%) |
| Leu (L) | 30 (7.7%) | 19 (6.4%) | 20 (6.7%) | 25 (6.4%) | 20 (7.2%) | 26 (8.9%) | 20 (6.6%) |
| Lys (K) | 26 (6.6%) | 17 (5.7%) | 14 (4.7%) | 24 (6.2%) | 22 (7.9%) | 20 (6.8%) | 25 (8.3%) |
| Met (M) | 12 (3.1%) | 6 (2.0%) | 9 (3.0%) | 12 (3.1%) | 10 (3.6%) | 7 (2.4%) | 7 (2.3%) |
| Phe (F) | 19 (4.8%) | 17 (5.7%) | 17 (5.7%) | 19 (4.9%) | 15 (5.4%) | 18 (6.1%) | 17 (5.6%) |
| Pro (P) | 13 (3.3%) | 14 (4.7%) | 17 (5.7%) | 16 (4.1%) | 7 (2.5%) | 10 (3.4%) | 12 (4.0%) |
| Ser (S) | 42 (10.7%) | 28 (9.4%) | 28 (9.4%) | 35 (9.0%) | 26 (9.4%) | 18 (6.1%) | 21 (6.9%) |
| Thr (T) | 22 (5.6%) | 13 (4.3%) | 10 (3.4%) | 23 (5.9%) | 16 (5.8%) | 16 (5.5%) | 15 (5.0%) |
| Trp (W) | 2 (0.5%) | 1 (0.3%) | 1 (0.3%) | 2 (0.5%) | 2 (0.7%) | 2 (0.7%) | 2 (0.7%) |
| Tyr (Y) | 16 (4.1%) | 9 (3.0%) | 9 (3.0%) | 15 (3.8%) | 10 (3.6%) | 10 (3.4%) | 9 (3.0%) |
| Val (V) | 20 (5.1%) | 23 (7.7%) | 24 (8.1%) | 24 (6.2%) | 13 (4.7%) | 18 (6.1%) | 18 (5.9%) |
| Molecular Weight (Da) | 44192.61 | 33437.75 | 33192.51 | 43727.11 | 31,608 | 33,310 | 34,277 |
| pI | 5.75 | 5.87 | 5.42 | 5.49 | 6.09 | 6.46 | 5.72 |
| Extinction Coefficient (280 nm, $M^{-1}cm^{-1}$) (2) | 34840 | 18910 | 18910 | 33350 | 26,500 | 25,900 | 24,410 |

1 Parameters exclude the LEHHHHHH C-terminal tag.

2 Assuming all cysteine residues are reduced.

**Supplementary Table S4:** PaCasp7a crystal statistics.

| | | | |
|---|---|---|---|
| **PDB Code** | | **Refinement** | |
| **Data collection** | | $R_{work}/R_{free}$ **(%)** | 17.7/21 |
| **Wavelength** | 1.0 | **Average B-factor (Å$^2$)** | 17.76 |
| **Temperature(K)** | 100 | **Macromolecules** | 16.40 |
| **Space Group** | P $2_1$ $2_1$ $2_1$ | **Solvent** | 27.77 |
| **Cell Dimensions** | | **Wilson B-factor** | 13.39 |
| **a, b, c (Å)** | 74.416<br>86.848<br>93.666 | **R. m. s. deviations** | |
| **α, β, γ (°)** | 90.00<br>90.00<br>90.00 | **Bond length (Å)** | 0.077 |
| **#Unique Reflection** | 83910 | **Bond angle (°)** | 5.004 |
| **Resolution (Å)** | 39.4 – 1.57 | **MolProbity score** | 4.84 |
| **R-meas** | 0.249 | **Number of atoms** | |

| | | | |
|---|---|---|---|
| **R-pim** | 0.117 | **Protein** | 3823 |
| **CC(1/2)** | 0.255 | **Water** | 586 |
| **Average I/σ** | 1.33 | **Protein residues** | 466 |
| **Completeness (%)** | 98.5 | **MolProbity** | |
| **Redundancy** | 4.9 | **Ramachandran favored** | 97.86% |
| **Clash score** | 3.42 | **Ramachandran outliers** | 0.00% |
| | | **Rotamer outliers** | 0.24% |

**Supplementary Table S5:** Molecular weight determined using ProtParam and mass spectrometry.

| Protein | M.W. Full length (kDa) (Protparam) | M.W. Large Subunit (kDa) (Mass Spec.) | M.W. Small Subunit (kDa) (Mass Spec.) | M. W. A Protomer (cleaved IL and prodomain) |
|---|---|---|---|---|
| OfCasp3a | 45.3 | 17.9 | 13.2 | 31.1 |
| OfCasp3b | 34.5 | 20.3 | 12.1 | 32.4 |
| PaCasp3 | 34.3 | 18.9 | 12.9 | 31.8 |
| PaCasp7a | 44.8 | 17.8 | 13.3 | 31.1 |

**Supplementary Table S6**: Homologs of components of human apoptotic pathways in *O. faveolata* and *P. astreoides*.

| Protein_name | Uniprot ID (Human) | *Orbicella faveolata* | *Porites astreoides* |
|---|---|---|---|
| Bcl-2 | P10415 | XP_020601884.1 | comp77460_c1_seq5.p1 |
| Bcl-xL | Q07817 | XP_020600881.1 | comp71697_c0_seq4.p1 |
| Bax | Q07812 | XP_020620765.1 | comp65965_c0_seq1.p1 |
| Bak | Q16611 | XP_020601942.1 | comp78640_c3_seq2.p1 |
| PIDD | Q9HB75 | XP_020624054.1 | comp63514_c0_seq4.p1 |
| RAIDD | P78560 | XP_020606624.1 | comp74936_c0_seq2.p1 |
| P53 | P04637 | XP_020628162.1 | comp71445_c0_seq1.p1 |
| Apaf-1 | O14727 | XP_020620792.1 | comp76833_c0_seq1.p1 |
| Cytochrome c | P99999 | XP_02062022.1 | Isotig14309.p1 |

```
OfCasp8a   1 MDSSAETTRLLYRISNEMRSQDLQNLKHLCHERIPLGELEKATQASALFQIMLQKRLIRPENLSFLEGLLNQIGRADLAG 80

OfCasp3a   1 ------------------------------------------------------------MEQSDRDILRKNRQDLLKDM 20
PaCasp7a   1 ----------------------------------------------------------------MQEEDRKALRSNREAL 16
PaCasp2    1 -------------------------------------------------------------MDKKHRDLLRKNRISLVQD 19
OfCasp2    1 -------------------------------------------------------MDKKHRELLRKNRLALVQDLEAAQL 25
OfCasp8a  81 KIQSSGRDVTMQTESALGSNSVDRNYRAFLMKLSDELTRDNVESLKFVANLPDGIEEKVQNGKDLFKYMIQQGNLGPCKL 160

OfCasp7    1 ---------MEKDECRSLSKSICIEFARDQVLRAQQLLERFRLCDCPDVGGFECRCVSEEVMETIGNRSYTERSTQTEEN 72
PaCasp7b   1 -----MEKDLCRTIVKRISTGYAHDRMQKAQQLLERIRFCDCMAFGGKRCACVRRPREVLNGRNNTRDKGTQTEERTGAT 75
PaCasp3    1 ------------------------------------------------------------------MSKFNTVVSGCAGN 14
OfCasp3c   1 -------------------------------------------------MEFEQWPGTLMVQNNDSLNIRSIPRYGHVPC 31
OfCasp3b   1 ---------------------------------------------------------------MSSKFKIHVADCSGAVV 17
OfCasp3a  21 EAKRVASRLYSRGIFSEEDKDEVNSKSTTNEQGECLLDILPKRGPKAFRAFCDVLHELSPHLESLLRPVQEAGIPSEGTD 100
PaCasp7a  17 LKDLEAKRVASLLYSREIFSEEDKDSVNAKSTPSEQREEVLDILPRKGPKAFSAFCNVLHEVSPHLEALLRPIQEDDEGI 96
PaCasp2   20 LEATQLLSYLYQEDILSENDLDSIKAEKTRGGKAEKLLDDILPRRGKKAFDVFCQALATTDGQGHLVDLLKPNESISSGN 99
OfCasp2   26 LNYLFQENCLSENDVDSIKVQATPRAKAEKLLDILPRRGPQSFDVFCRALANTDGQGHLVDLLRTNESIASGAISSTTE 105
OfCasp8a 161 DHLADMLQEIHRRDLAEKVKKFQSEGGVSPSTSDDQPPPMQINNQRQMVVTPPSGRPLVTGKQPTMHQMGESRPPAVPLL 240
OfCasp8b   1 ---------------------------------------------MIVLFTQTFPLQSLVITFPTESKEEKKKSRRIVEK 35

OfCasp7   73 NRSTPLKSSLFYRSDSSSAADLRDGPPNFNIQRNVVSVARNYFPDDQTYDGIKNP--YVLVINNVNFYNDPR----PRNG 146
PaCasp7b  76 TAQYFGHLDREDAANPTYAVPQPTISLLDVNRNVVPLAKRRHLPEDDVYDGIRNP--YVLIINNVNFWYIPQ----PRTG 149
PaCasp3   15 IVVGDHAVLNVGASSTDARPRKQPPSEEVPPTGPSQVHCQESAGDGMNYIAK---SGYVLVINNYLFPQRLNVE---RTG 88
OfCasp3c  32 HGHNNNYKAGTWQFSGGNNNYKAGTWQFSGGNNNYKAGTWEFSGDNNNYKAK---SGYVLIMNNYIFPKRDDVE---RIA 105
OfCasp3b  18 IGDHANLTVEAPVAGAGPQRRPVREQAAAATTTSTHAPCQESSGDDINYKAK---SGYVLVINNYIFPRRDDVE---RTG 91
OfCasp3a 101 GGNKNPISSTSNLGASESADEADAKLFGFGGGSASAKPSSSTLDKEIIYKMDRSTRGIAVIINNKNFLRSSGMDRYPRNG 180
PaCasp7a  97 DGPGSTAPIPASGVPSENNDQADAKLFSFGGGGSAAKSSANTIDNDTIYKMNKSTRGIAVIINNKDFLRSSGMDRYPRNG 176
PaCasp2  100 VSSTTAGNAVEVNDGVANTAHHVAVKSNVETSSQVKKNVLPGTDEDFVYPMNCKPHGWCLIVNNVDFETLRR-----RGG 174
OfCasp2  106 NIGKISSEMNDGNSSPFPSHQVKVVKPNVSFSSQVKKNVSGSTDEDFVYSMKCKPHGWCLIVNNVDFEHLNR-----RGG 180
OfCasp8a 241 SPSFFSRAQTSDAGNVEPRYPVQAMPNPAEILAPMQQLGVDEEEAMPSYPMTNRPRGIALIISNEHFQGNPDLR--DRQG 318
OfCasp8b  36 LKGELEWLQRDNERLEEDNKIFKQREEEREAEIERFQRDKEHATELDRYEMEQGRRGICLVINNYNFPRGRE-----RQG 110

OfCasp7  147 ARHDLDNLKRFMKEAGFRVRIHSDLRKKQMLAILEETRLNGALAEHDSFICIIMSHGNEEGILGIDSQAIPVDDITAKFQ 226
PaCasp7b 150 AQHDFDNVMSFVKKAGFRVVQCPDLRKSKLLDLLEETRQSEVLVEHDSFICFIMSHGNEEGILGVDSRAIPVDEITAKFQ 229
PaCasp3   89 SNDDVKNLTNLFDDFGFRSRVQDNQTRSEMLSLLKDTAEK-DFSKYDCFVCVILSHGSKDGIYGTDDEVINIEAITSLFR 167
OfCasp3c 106 SNDDVRNLTSLFDDFNFRHRLENNQTGSQMINLLQTTAEK-DFSGYDCFVCVILSHGGKDGIYGTDEELIKIEAITSLFR 184
OfCasp3b  92 SNDDVRNLTSLFDDFNFRHVVEDNQTGSQMIKLLQTTAEK-DFSRYDCFVCVILSHGKKDGIYGTDEELIKIEAITSLFR 170
OfCasp3a 181 TDVDRDSLVKLFRMLKFEVKIYNDRTKAEIRTITKEMATL-NHSNYDAFIFSILTHGEEGVIYGTDG-TISIRDLTAEFK 259
PaCasp7a 177 TDVDRDALAKLFRALKFDVRIYNNQTRAEIRRITKEMAIT-NHTPYDAFIFSILTHGEEGVIYGTDG-TMAIKDLTAIFK 255
PaCasp2  175 SDLDAKQLSELFTQLHYKVKVVRNQTAKQLKDTLFQFAKMSDHKAADSMVLCLLSHGLEGQIYGVDGILVSIPDLLALFN 254
OfCasp2  181 SDLDAQKLEELFTQLQYKVKMFRNQTSKQLKETLLQFAKLDDHRVADSMVVCLLSHGLEGQIYGIDGVLVSIPDLLALFN 260
OfCasp8a 319 NEKDVKHLRELWEFLNFEVIVRADLESEDIYDVLRGISTM-DHRNFDCFVCCLLSHGANGGIYGTDSTLVEIKDITSMFK 397
OfCasp8b 111 AEIDEKKIVDLFRDLDFEVKFLQNLLGSEILKAAKEFAAR-DLSQFTEFVFIIMSHGDKDAIEGVDGAFVKVAPLMSLFS 189

OfCasp7  227 GGQCPQLVTKPKLFFLQACRGDVDDKGYYVPRGPRSQDVY-------------ADSDHFQEMPVKLPSDADFLIAYSTTK 293
PaCasp7b 230 GGKCPYLATKPKLFFLQACRGDVDDEGYLVPRGPGHVY---------------PDA--IDEMPVKLPSDADFLIAYSTTK 292
PaCasp3  168 RDECPSLEGKPKIFLIQACRGNQRDIVPVESDSDP-----------------------MVFSNPSLPADADFLICFASAP 224
OfCasp3c 185 CNECPSLEGKPKIFLIQACRGSQRDRVTVESDGPA-----------------------ILPSWSSSPADADFLICFASAP 241
OfCasp3b 171 RNECPSLEGKPKIFLFQACRGSRRDTVPVESDSDP-----------------------IPLSSSSLPADADFLICFASAP 227
OfCasp3a 260 Y--STSLAGKPKLFFFQACQGHEYMDGMDVTDAP-------------------------QENKVSVPAEADFLYAYSTVP 312
PaCasp7a 256 D--CTTLVGKPKMFFFQACQGHEYMDGMDVTDAP------------------------QQGSKVSVPAEADFVYAYSTVP 309
PaCasp2  255 GYMAKDLIGKPKLFFIQACRGSDYDHGTDMTDGGHSSNVEKEYHVTRSTNELLAAACPNDKPETTIPAEADMLVAYSTVP 334
OfCasp2  261 GYMAKDLVGKPKLFFIQACRGSDFDHGADMTDGAHIPGDLHVKQS---TEQLLNAAFPTD----TLPAEADTLVAYSTVP 333
OfCasp8a 398 GVACPSLAHKPKLFFIQACRGLDRDRGARFETDAVESN------------------PEDAMRHNAEPNESHFLLGYATPP 459
OfCasp8b 190 VTKCPTLRNKPKLFFIEACRGPRKDRLPARLRNTEGNGIA-------------MMLCADSNLAHGVPLEADFLLSFATVP 256

OfCasp7  294 GYVSHRRFIDGSWFISCLVQVLRANSHKEDLMTMLTRVNKAM-SEFYTE------GGSKQISCQLSMLTKKVYFASFLNK 365
PaCasp7b 293 GHLSNRRFADGSWFISCLIQILETYSGREDLMAMLTRVNRSMCKEYSVP------GGFKQISCQLSMLTKKVYFTNFQ-- 364
PaCasp3  225 GHQSYRQPLLGSWFISAVFDVFKEHAEREHIMDMMLRVNNQV-AGYYSR------DGLKQMPCQICMLTKKVFFQPRYGS 297
OfCasp3c 242 GHESYRQVGIGSWFIAAVVDVFKEYAEREHLMDMMTRVNDRV-AGFFSK------EGLKQMPSQVSMLRKKVFFDPKYS- 313
OfCasp3b 228 DHESYRQAHLGSWFISSVVDVFKEYAEREHLMDMMLRVNNHV-AGFYSK------EGLKQIPCQVCMLRKKVFFDPKYS- 300
OfCasp3a 313 GYYSWRNSVNGSWFIQSLTKVFEDNAERMDILRMLTRVNAMV-STYKSRTGDYYSDSKRQVSSIVSMLRKELYFFPENVE 390
PaCasp7a 310 GYYSWRNSVNGSWFIQSLTKVFEENAERMDILRMLTRVNAMV-STYKSRTGDYYSDSKRQVSSVVSMLRKELYFFPENVT 387
PaCasp2  335 GYVSWSNLQKGSWFVQALVDVFSAYAKAEDVVSMLVKVNGKV-AREFESF-----NRKKQMPAPVIMLTKKVFFFAQS-- 405
OfCasp2  334 GYVSWSNLQKGSWFVQAIVDVFSAYAHEEDVVSMLVRVNGKV-AREFESF-----NRKKQIPAPVIMLTKKVFFFPS--- 404
OfCasp8a 460 GYVSWRSQVHGSWYVSKLCEVFRKYCERFDVMTMMVKVNDEV-SDAFTR------QGYKQCPAPVVTLRKRIYLNNN--- 529
OfCasp8b 257 GYKAWRSQVAGSWYIQVLADVIRKYHRHHHLLDMLTEVNKQV-GAIANDYP--EHDLSFQVPAQSNTLRAKVFL------ 327
```

**Supplementary Figure S1:** Multiple sequence alignment of caspases from *O. faveolata* and *P. astreoides*.

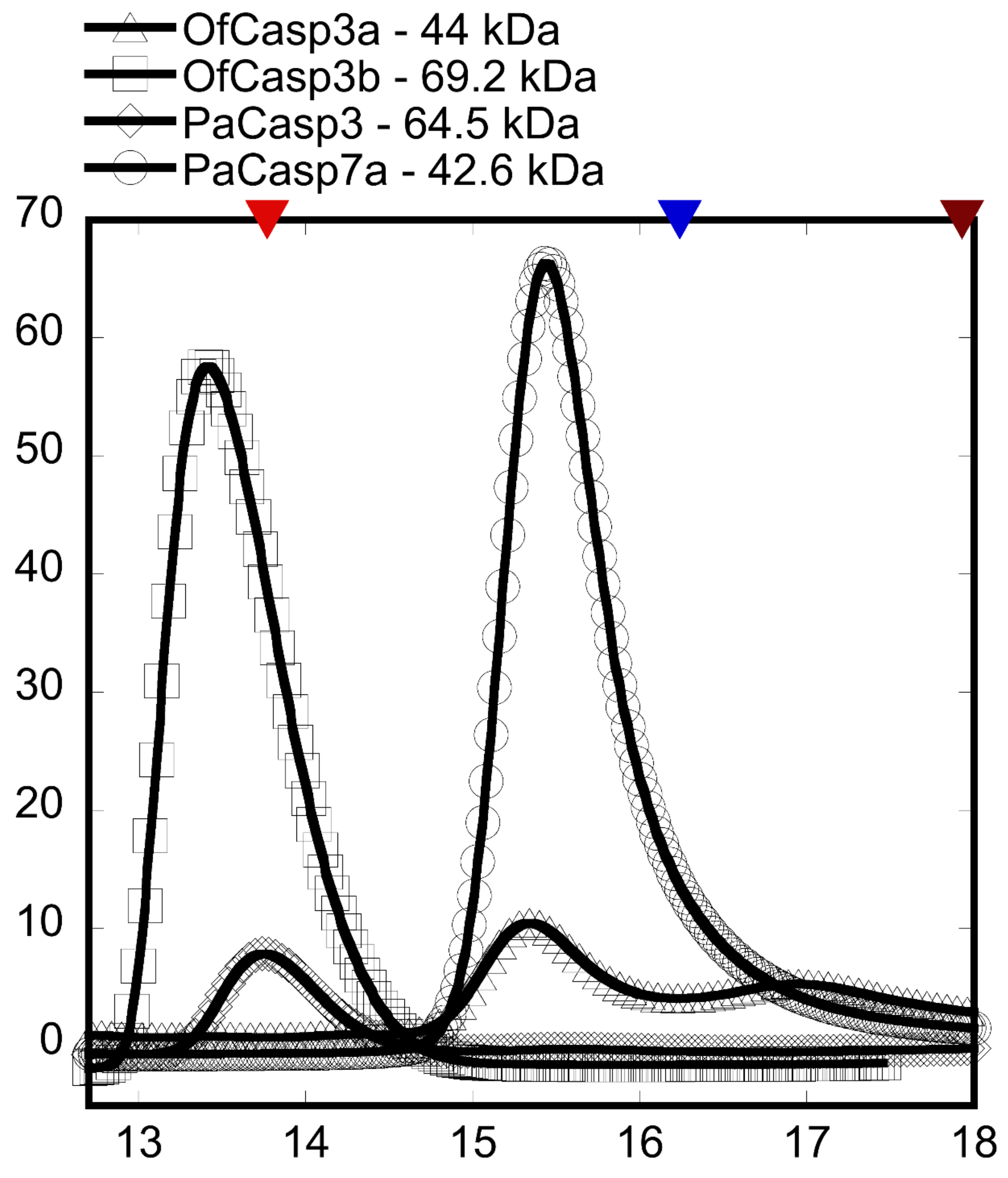


**Supplementary Figure S2:** Chromatogram ($A_{280}$) of coral caspases. Peaks based on elution volume after FPLC analysis of the native oligomeric state through sizing column. Sigma-Aldrich gel filtration kit was used as a marker; Albumin (66 kDa), Carbonic anhydrase (29 kDa) and cytochrome c (12.4 kDa).

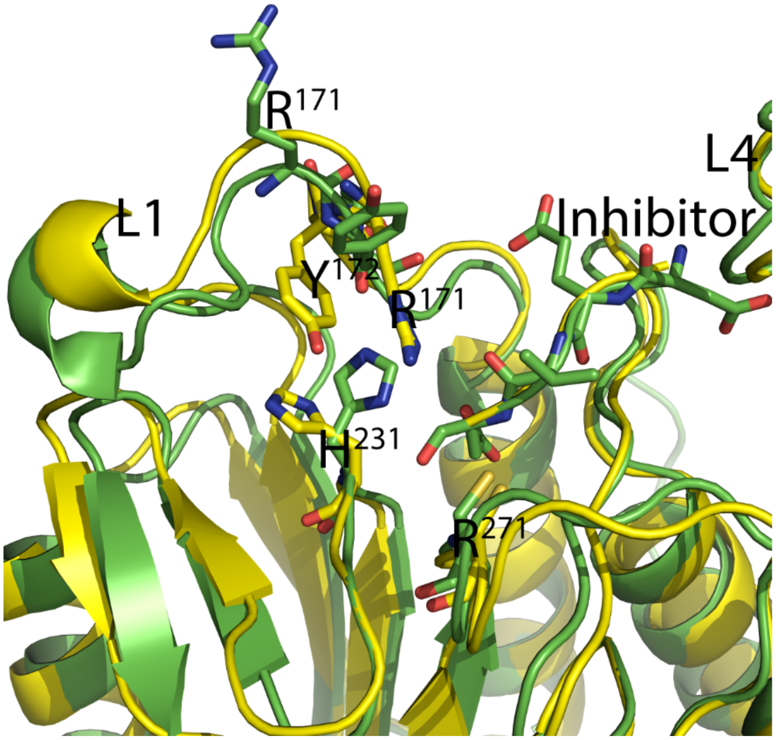


**Supplementary Figure S3:** Average structures of molecular dynamics (MD) simulations showing loop 1 (L1) containing "RYP" motif in the "in" *versus* "out" conformations. Green: Past7a crystal structure and "RYP out", Yellow: PaCasp7a average structure from MD simulations of PaCasp7a model structure and "RYP in". The data show that the two conformations do not interconvert on the timescale of the MD simulations (50 ns).

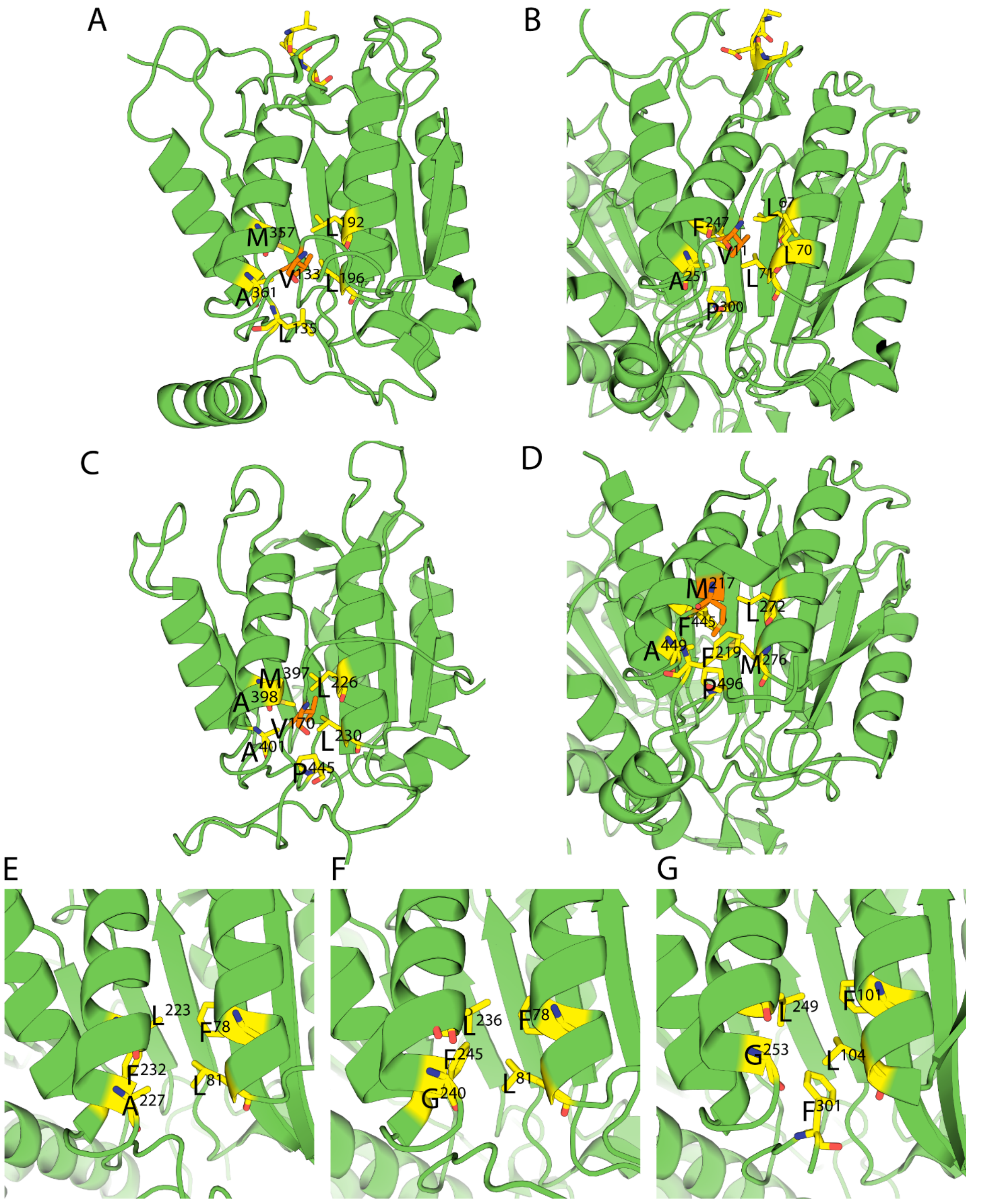


**Supplementary Figure S4:** Structures of caspases with N-terminal peptide bound between helices 1 and 4. A. HsCasp1 (PDB ID: 2H48), B. HsCasp2 (PDB ID: 3RJM), C. *D. melanogaster* initiator caspase Dronc (PDB ID: 2FP3), D. *C. elegans* caspase CED-3 (PDB ID: 4M9R), E-G. Hydrophobic pocket between helices 1 and 4 in human effector caspases. E. HsCasp3 (PDB ID: 2J30), F. HsCasp6 (PDB ID: 3S70), G. HsCasp7 (PDB ID: 1F1J).
